# N153-linked glycans on envelope protein protect orthoflaviviruses from antibody-mediated clearance

**DOI:** 10.1101/2025.02.20.639387

**Authors:** Donald Heng Rong Ting, Jan Kazimierz Marzinek, Corrine Wan, Raghuvamsi V. Palur, Fakhriedzwan Idris, Eunice Tze Xin Tan, Wei Teng Clara Koh, Xin Ni Lim, Tun-Linn Thein, Cheer Vee Lim, Chin Huan Ng, Theresa S. C. Buckley, Ian Walsh, Paul A. MacAry, Yee-Sin Leo, Ganesh S. Anand, Kuan Rong Chan, Terry Nguyen-Khuong, Peter John Bond, Sylvie Alonso

## Abstract

The envelope (E) protein of dengue virus (DENV) is glycosylated at two highly conserved asparagine (N) sites (N67 and N153). The role and importance of these N-linked glycans in DENV pathogenesis has been elusive. Here, we report the critical role of N153-linked glycans on E protein in preventing antibody-mediated viral clearance. A DENV2 mutant lacking N153-linked glycans (N153Q mutant) was engineered and found to be mildly impaired *in vitro* but drastically attenuated in a symptomatic mouse model of severe dengue, as evidenced by accelerated viral clearance. In B cell-deficient mouse models, N153Q mutant displayed parental virulence and viremia profile. Homologous and heterologous passive transfers of purified IgM from infected B cell-proficient mice into B cell-deficient mice demonstrated the role of N153Q-specific IgM in N153Q attenuation and accelerated clearance, while WT DENV was unaffected by IgM from both WT- and N153Q-infected mice. Furthermore, *in vitro* neutralization assay supported that the accelerated clearance of N153Q mutant in mice was mediated by non-neutralizing IgM. Furthermore, using plasma samples from convalescent dengue patients and monoclonal antibodies, *in vitro* neutralization assays showed that N153Q virus was more susceptible than WT to IgG-mediated neutralization. Glycoproteomics combined with molecular dynamics (MD) simulations revealed that glycan composition on E protein influenced IgG binding. Our findings were extended to all DENV serotypes and ZIKV, hence supporting that the N153 glycans-mediated immune evasion strategy is conserved across orthoflaviviruses.

## Introduction

Dengue is a highly prevalent mosquito-borne disease that poses a significant burden on affected societies, primarily in more than 120 countries within the tropical and subtropical belt [1]. It is estimated that there are 390 million annual infections, and around 6.1 billion people will be at risk of contracting the disease in 2080 [2, 3]. The disease is caused by four distinct serotypes of dengue virus (DENV). Individuals infected with any one of these serotypes can display a wide spectrum of clinical manifestations that have been classified by the World Health Organization (WHO) as dengue with or without warning signs, and severe dengue [4]. Mild cases manifest as skin rashes, fever, headache, joint and muscle pain, nausea and vomiting, and mild bleeding. In severe cases, patients experience plasma leakage, hemorrhage and organ impairment, which can be fatal if not carefully managed [4]. The increased risk of developing severe disease upon secondary infection with a heterologous serotype has been linked to antibody dependent enhancement (ADE), where binding but non-neutralizing antibodies enhance virus infection [5, 6]. Despite being a serious public health concern, there is currently no effective antiviral treatment, while the development of one of the most promising small molecule inhibitors, JNJ-1802 (Johnson & Johnson) that had entered the clinical pipeline [7–10], has recently been discontinued. There are also two licensed tetravalent live attenuated vaccines, namely Dengvaxia® (Sanofi Pasteur) and QDENGA® (Takeda’s Biologics). However, both vaccines confer serotype-dependent protective efficacy, which greatly limits the impact of vaccination in the majority of countries where more than one DENV serotype co-circulate with unpredictable dominance patterns [11–14]. Furthermore, Dengvaxia® was found to increase the risk of severe disease in seronegative populations, and hence its use has been limited to individuals with pre-existing DENV immunity [15].

DENV is a positive single-stranded RNA enveloped virus that belongs to the Orthoflavivirus genus [16]. Its 11 kb viral RNA genome encodes three structural proteins (capsid [C], pre-membrane [prM], and envelope [E]) and seven non-structural proteins (NS1, NS2A, NS2B, NS3, NS4A, NS4B, and NS5). The main viral structural protein, E protein, is a vaccine candidate of choice currently being tested in clinical trials [17]. E protein consists of three domains each with distinct functions, and harbors two highly conserved N-linked glycosylation consensus motifs (NXS/T) at N67 and N153, which are located on domain II and domain I respectively [18]. No O-linked glycosylation has been reported on E protein [19]. Both N67- and N153-linked glycosylation sites are conserved in all DENV strains from the four serotypes. N153/N154-linked glycosylation site is also found in other closely related orthoflaviviruses such as Zika virus (ZIKV) and West Nile virus (WNV), while N67-linked glycosylation site is unique to DENV.

Relying entirely on the host cell glycosylation machinery, glycoforms present at N67 and N153 sites on E protein of DENV were found to be highly heterogenous and dependent on the serotype and host cell type used in the study [20, 21]. Earlier studies have shown that N-linked glycans on E protein were involved in DENV binding to several receptors, including liver/lymph node– specific ICAM-3–grabbing nonintegrin (L-SIGN; also known as DC-SIGNR, CD209L), C-type lectin domain family 5 member A (CLEC5A; also known as CLECSF5 and MDL-1) and mannose receptor (MR), but these studies could not distinguish between both N-sites due to some limitations in the experimental approaches [22–24]. A structural study showed that N67-linked glycans interacted with C-type lectin dendritic cell-specific ICAM3 grabbing nonintegrin (DC-SIGN), which is one of the main DENV receptors [25]. Abolishment of the N67-linked glycosylation site resulted in comparable, reduced or completely abrogated growth in mosquito and mammalian cell lines, depending on the studies, thus making the role of N67-glycans controversial [26–28]. Absence of N153-linked glycans was shown to expose the fusion peptide on E protein, resulting in elevated pH threshold for endosomal membrane fusion, which in turn negatively impacted the ability of the virus to uncoat and release its RNA genome into the cytoplasm [29, 30]. N153 deglycosylated DENV mutants were able to replicate in both mammalian and mosquito cell lines but produced lower virus titers than their wildtype (WT) counterpart [26–28]. Furthermore, N153-linked glycans were proposed to cause steric hindrance and interfere with antibodies binding to the surface of the viral particle, although direct evidence is lacking; alternatively, or in addition, N153-linked glycans were shown to be the target of antibodies, leading to virus neutralization [31–34].

While no *in vivo* studies have specifically examined the role of N153-linked glycans of E protein in DENV, studies with other orthoflaviviruses (e.g., ZIKV, JEV, WNV) suggested that the absence of N153/N154-linked glycans impaired neurovirulence, reduced infectivity in DC-SIGN/L-SIGN bearing cells and triggered heightened innate cytokine responses [35–39].

Here, we investigated the role of N-linked glycans on E protein from a DENV2 Cosmopolitan strain, namely D2Y98P. DENV2 Cosmopolitan is the most widespread DENV2 genotype that has been circulating in Asia-Pacific, Middle East, and Africa for decades and has caused recurrent outbreaks associated with severe dengue [40]. More recently, Cosmopolitan DENV2 has expanded further into other parts of the world like South America [41], underscoring the exceptional fitness of these strains. Leveraging on the unique ability of this non-mouse-adapted virus strain to induce symptomatic, clinically relevant disease upon primary infection in Type I interferon-deficient mice (IFNAR^-/-^) mice [42], we uncovered the critical role of N153-linked glycans in preventing antibody-mediated viral clearance, representing a novel immune evasion strategy for DENV2. Our findings were extended to the other DENV serotypes and ZIKV, thus suggesting that this immune evasion strategy is widely shared across the orthoflavivirus genus.

## Results

### Generation of N67- and N153-deglycosylated DENV mutants

DENV2 mutants that lack glycan motifs at N67 or N153 on E protein were generated by substituting either the first or third amino acid of the N-linked glycosylation motif (N-X-T/S, with X being any amino acid except proline). N153-deglycosylated mutant (N153Q mutant) was successfully obtained and remained genetically stable after several passages in cell culture. In contrast, attempts to generate N67-deglycosylated mutants were unsuccessful, as these mutants rapidly regained the N-linked glycosylation motif upon passaging in cell culture (Suppl. Table S1). Furthermore, comparable deuterium uptake profiles were obtained between WT and N153Q E proteins (Suppl. Fig. S1), thus supporting that the N153Q amino acid substitution on E protein and lack of glycans at this position did not significantly alter the overall virion breathing dynamics as measured by HDXMS.

### N153Q mutant is slightly impaired in cell lines but is drastically attenuated in mice

The *in vitro* fitness of N153Q mutant was examined in mosquito and mammalian cell lines by monitoring the viral titers in culture supernatants. N153Q mutant retained parental *in vitro* fitness in C6/36 and BHK-21 cell line but was mildly attenuated in Vero and Huh-7 cell lines, with virus titers fold-reduction ranging between 2 to 8.4 compared to WT (Fig. 1A). We next determined the *in vivo* phenotype of N153Q mutant in Type I interferon receptor-deficient (IFNAR^-/-^) mice. All N153Q mutant-infected mice survived the infection, while only 10% of WT-infected mice survived (Fig. 1B). Infection with either strain led to significant body weight loss during the initial phase of infection (day 1-5 p.i). However, N153Q mutant-infected mice regained their body weight afterwards, while WT-infected mice kept losing weight and had to be euthanized (Fig. 1C). N153Q mutant-infected mice displayed transient, mild clinical signs (ruffled fur and hunched back), in contrast to severe symptoms (diarrhea and lethargy) displayed by WT-infected mice (Fig. 1D). Notably, N153Q mutant was rapidly cleared from the blood circulation, with viremia titers falling below the detection limit from day 3 pi. onwards whereas viremia titers in WT-infected mice kept increasing over time (Fig. 1E). Interestingly, both infected groups had comparable viral loads in perfused organs including spleen, axillary and branchial lymph nodes throughout the course of infection (Fig. 1E). In contrast N153Q mutant was barely detectable in perfused kidneys and liver, whereas WT virus titers were detected at day 3 and 4 pi. (Fig. 1E).

**Figure 1.**
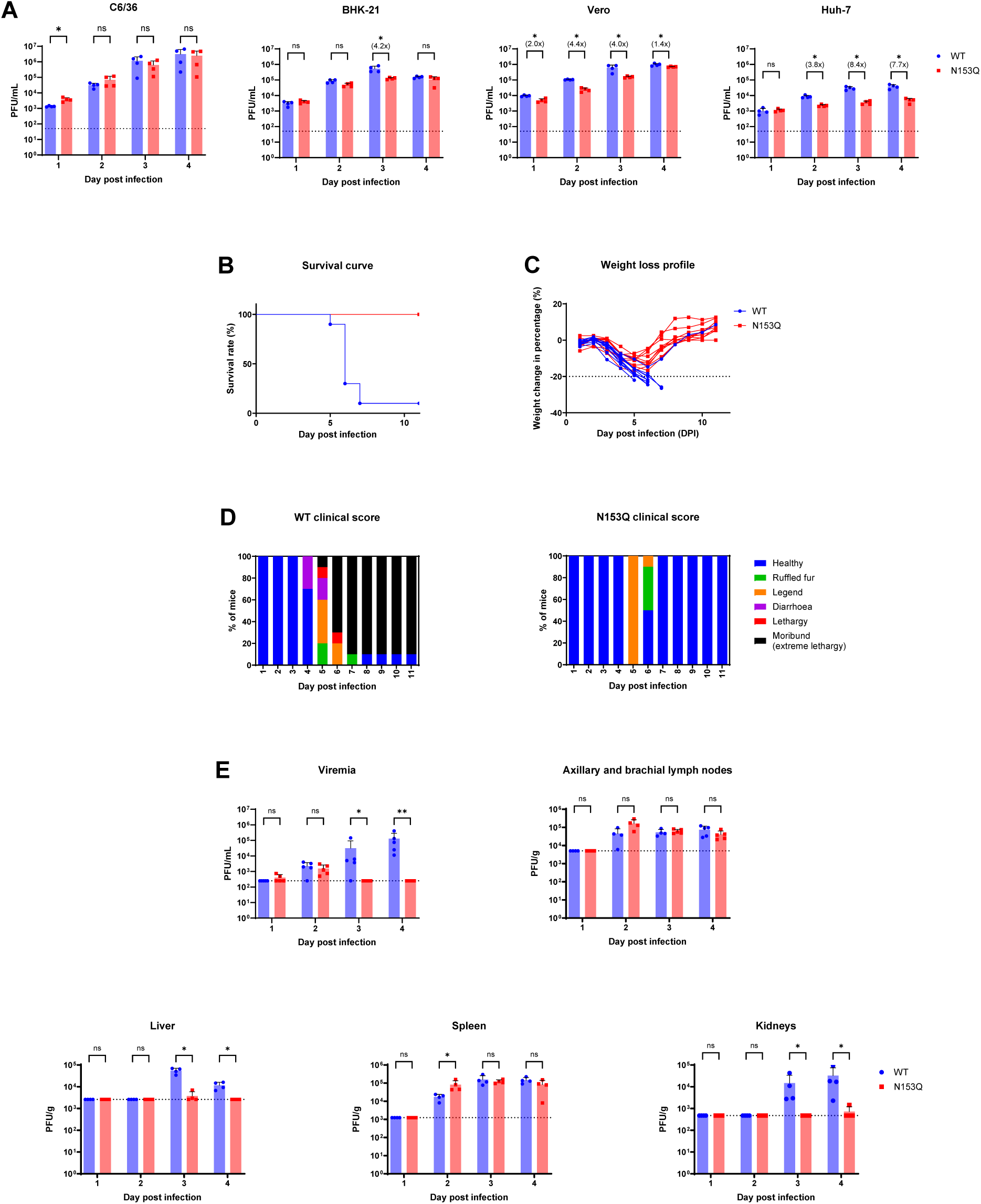
*In vitro* and *in vivo* characterization of N153Q DENV2 mutant. (A) Cells were infected with WT or N153Q DENV2 at MOI 0.1. Viral titers in supernatant were measured by plaque assay (n=4). (B-E) 5-7 weeks old IFNAR^-/-^ mice were sc. infected with WT or N153Q mutant at 10^5^ PFU/mouse. (B) Kaplan-Meier survival curve (n=10). (C) Body weight loss profile (n=10). (C) Clinical scores (n=10). Mice that lost 20% of their body weight measured at the start of infection and/or that were moribund were promptly euthanized. (E) Viremia titers and viral loads in perfused axillary and brachial lymph nodes, liver, spleen and kidneys (n=4-5) were measured by plaque assay and normalized against wet organ weight. Dotted line indicates the detection limit of the assay. Data shown are representative of at least two independent experiments. Statistical analysis was performed using nonparametric Mann–Whitney U test. *, p < 0.05; **, p < 0.01; ns, not significant.

Together, these results indicated that N153-linked glycans on E protein were largely dispensable for *in vitro* viral fitness but played a critical role in viral fitness and virulence in IFNAR^-/-^ mice. In this symptomatic mouse model, N153Q mutant underwent accelerated clearance from the blood circulation and displayed impaired tropism in selected organs.

### N153Q mutant attenuation in mice is not due to differential host responses but involves B cells

To further investigate the *in vivo* attenuation of N153Q mutant, several blood parameters including the total and relative number of red blood cells (RBC), white blood cells (WBC) and platelets, as well as morphological changes of RBC and platelets, were determined in WT- and N153Q mutant-infected IFNAR^-/-^ mice from day 2 to 4 pi. However, no major differences were observed between both infected groups (Suppl. Fig. S2). Next, we examined the transcriptomic profile in WBC harvested at day 2 pi., time point at which comparable viremia titers were measured in both infected groups. Here again, there were no major differences in the transcriptomic profiles between both infected groups, with only 7 differentially expressed genes (DEGs) which displayed less than 3-fold change in their expression (Fig. 2A & Suppl. Table S2). Finally, we measured the systemic concentration of a panel of inflammatory cytokines. Elevated levels of TNFα, MCP-1, IL-10, IL-6, IL-27, IL-17A, IFNγ, and GM-CSF were detected in WT- and N153Q mutant-infected mice, compared to the uninfected controls (Fig. 2B & Suppl. Fig. S3). However, both infected groups displayed comparable levels.

**Figure 2.**
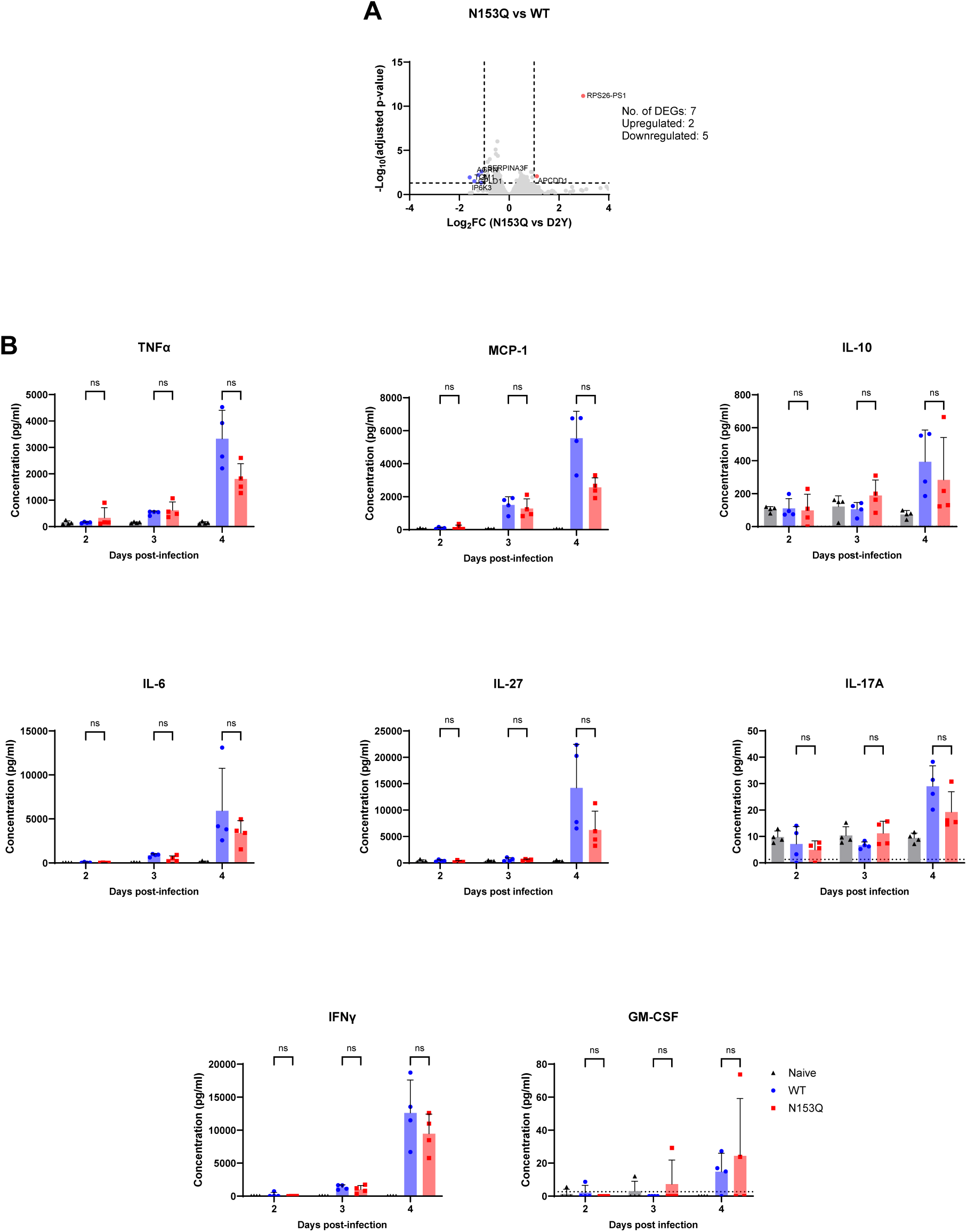
Transcriptomic and systemic inflammatory cytokines profiles in IFNAR^-/-^ mice infected with WT or N153Q mutant. (A&B) 5-7 weeks old mice (n=3 or n=4) were sc. infected with WT or N153Q DENV2 at 10^5^ PFU/mouse. (A) Total RNA was extracted from white blood cells of infected mice at day 2 pi. and subjected to bulk RNAseq. Volcano plot of differentially expressed genes between N153Q vs WT, where log2 fold-change of 1 and p-value < 0.05 was used as cutoff to the pairwise analyses. (B) Systemic concentrations of TNFα, MCP-1, IL-10, IL-6, IL-27, IL-17A, IFNγ and GM-CSF were measured by multiplex assay at the indicated timepoints post-infection. Data are representative of at least two independent experiments. Statistical analysis was performed using nonparametric Mann–Whitney U test. *, p < 0.05; ns, not significant.

Together, these results supported that N153Q mutant attenuation in IFNAR^-/-^ mice was not due to differential host responses to the infection.

The lack of differential innate immune responses to infection between WT and N153Q mutant in IFNAR^-/-^ mice prompted us to examine the role of antibodies in the accelerated clearance of N153Q mutant observed at day 3 pi. To do so, we evaluated the infection outcome and viral titers in IFNAR^-/-^ mice that were treated with an anti-CD20 monoclonal antibody (mAb), previously shown to deplete B cells [43, 44]. Treatment with anti-CD20 mAb significantly reduced anti-DENV IgM and IgG levels in WT-infected IFNAR^-/-^ mice, confirming the effective depletion of B cells (Suppl. Fig. S4A-B). Notably, N153Q mutant-infected IFNAR^-/-^ mice treated with anti-CD20 mAb had significantly higher viremia titers at day 3 and 4 pi. compared to the isotype mAb-treated group, with 17.1-fold and 114-fold increase respectively (Suppl. Fig. S4C). In contrast, B-cell depletion only resulted in moderate increase in WT viremia titers at day 3 and 4 pi. compared to the isotype mAb-treated control group, with 1.2-fold and 2.9-fold respectively (Suppl. Fig. S4C).

The involvement of B cells in N153Q mutant attenuation was further demonstrated in B cell-deficient IFNAR^-/-^ mice (muMT IFNAR^-/-^). In line with the observations made in CD20 mAb-treated IFNAR^-/-^ mice, infection of muMT IFNAR^-/-^ mice with N153Q mutant led to 100% mortality by day 7 pi. and was as lethal as infection with WT DENV (Fig. 3A), as evidenced by rapid body weight loss and severe clinical manifestations (Fig. 3B-C). Importantly, WT- and N153Q mutant-infected muMT IFNAR^-/-^ mice had comparable viremia titers, in sharp contrast to the rapid clearance of N153Q mutant seen in IFNAR^-/-^ mice (Fig. 3D). N153Q mutant maintained infection in the spleen, axillary and brachial lymph nodes in muMT IFNAR^-/-^ mice (Fig. 3D). However, lower N153Q viral titers were measured in the liver and kidneys (Fig. 3D).

**Figure 3.**
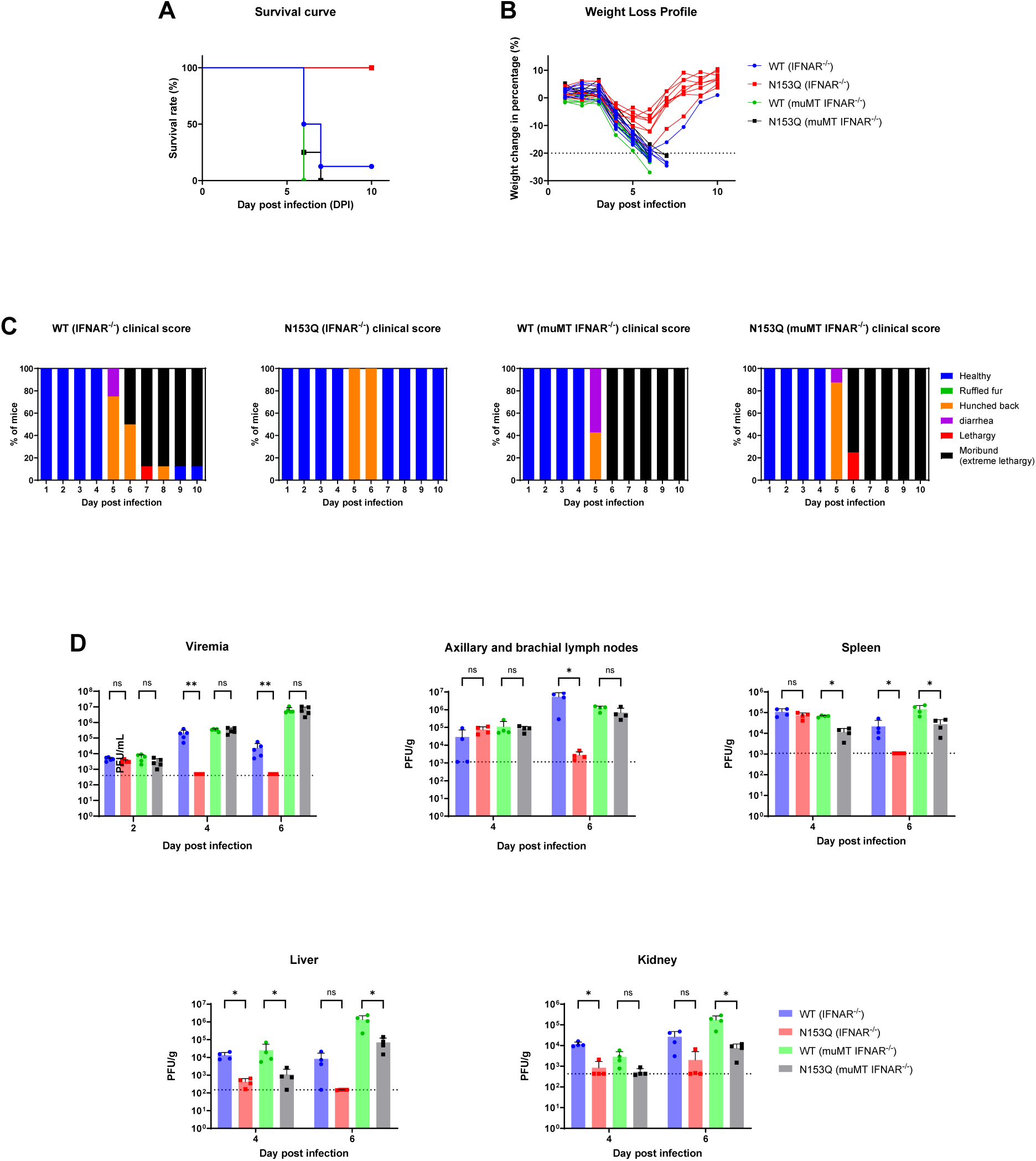
Infection profiles in muMT IFNAR^-/-^ mice. 5-7 weeks old IFNAR^-/-^ or muMT IFNAR^- /-^ mice (n=7/8) were sc. infected with WT or N153Q DENV2 at 10^5^ PFU/mouse. (A) Kaplan-Meier survival curve. (B) Body weight loss profile. (C) Clinical scores. Mice that lost 20% of their body weight measured at the start of infection and/or that were moribund were promptly euthanized. (D) Viremia titers and viral loads in perfused axillary and brachial lymph nodes, spleen, liver and kidneys (n=4-5) were measured by plaque assay and normalized against wet organ weight. Dotted line indicates the detection limit of the assay. Data are representative of at least two independent experiments. Statistical analysis was performed using nonparametric Mann–Whitney U test. *, p < 0.05; **, p < 0.01; ns, not significant.

Together, these results indicated that B cells played a critical role in the attenuated phenotype displayed by N153Q mutant in IFNAR^-/-^ mice. Restoration of parental viremia titers in B-cell deficient mouse models strongly supported the involvement of B cells (and antibodies) in the accelerated viral clearance of N153Q mutant in IFNAR^-/-^ mice. The lower viral titers measured in the liver and kidneys from N153Q mutant-infected muMT IFNAR^-/-^ mice further supported a role for N153-linked glycans in selective organ tropism.

To explore whether these findings could be extended to other orthoflaviviruses, we generated a ZIKV mutant that lacks N154-linked glycans on E protein (N154Q mutant). Infection of IFNAR^-/-^ and muMT IFNAR^-/-^ mice with WT and N154Q ZIKV phenocopied the results obtained with WT and N153Q DENV2 (Suppl. Fig. S5), thus strongly supporting the notion that N153/N154-linked glycans on E protein of DENV and ZIKV play a critical role in evading B cell-mediated viral clearance.

### N153Q mutant attenuation is mediated by non-neutralizing IgM antibodies

As B cells exert their effector functions mainly through antibodies, we examined the involvement of antibodies in the *in vivo* attenuation of N153Q mutant. To do so, a homologous passive transfer experiment was conducted where heat-inactivated serum harvested from WT- or N153Q mutant-infected IFNAR^-/-^ mice at day 5 pi. was administered to WT- or N153Q mutant-infected muMT IFNAR^-/-^ mice, respectively (Fig. 4A). Passive transfer of naïve serum to WT-infected or N153Q mutant-infected muMT IFNAR^-/-^ had no significant impact on the survival, body weight loss, clinical manifestations and viremia titers (Fig. 4B-E). Strikingly, passive transfer of N153Q mutant serum rescued N153Q mutant-infected muMT IFNAR^-/-^ mice, resulting in 75% survival rate (Fig. 4B). The protective effect was evidenced by transient body weight loss and milder clinical manifestations (ruffled fur and hunched back). Importantly, passive transfer of N153Q serum significantly reduced the viremia titers in N153Q mutant-infected muMT IFNAR^-/-^ mice at day 4 and 6 pi. compared to untreated mice and mice treated with naïve serum (Fig. 4E). Hence, these results supported that antibodies were involved in the accelerated viral clearance of N153Q mutant in IFNAR^-/-^ mice. Furthermore, very interestingly, passive transfer of WT serum did not protect WT-infected muMT IFNAR^-/-^ mice (Fig. 4B-D), and only moderately and transiently reduced the viremia titers at day 4 pi. compared to controls (Fig. 4E). These observations thus suggested that WT virus is resistant to antibody-mediated clearance, unlike N153Q mutant.

**Figure 4.**
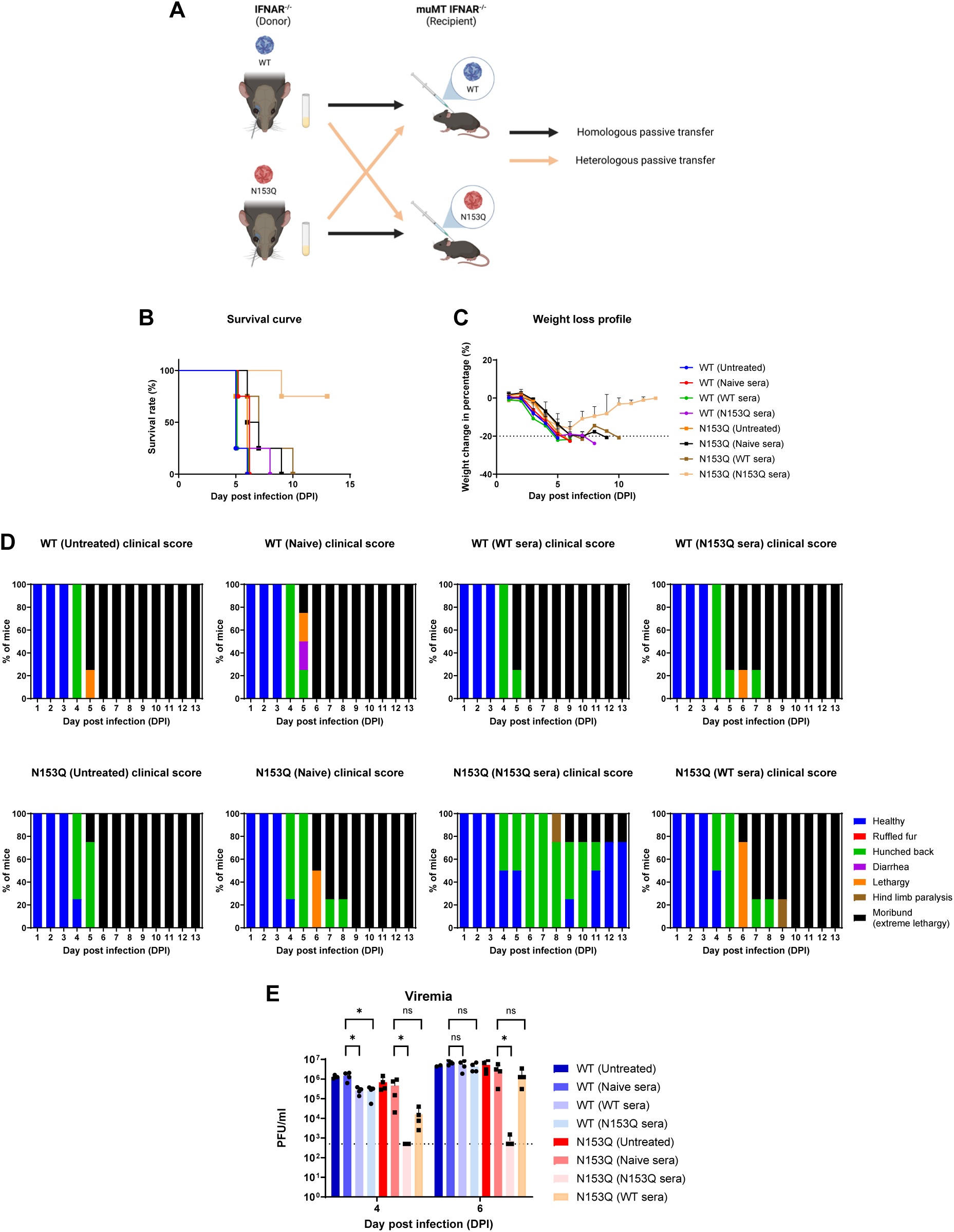
Homologous and heterologous passive transfer of serum in muMT IFNAR^-/-^ mice. (A) Schematic diagram of homologous and heterologous passive transfers. 5-7 weeks old muMT IFNAR^-/-^ mice (n=4) were sc. infected with WT or N153Q DENV2 at 10^5^ PFU/mouse. At day 3 pi. mice were administered intravenously with heat-inactivated serum from naïve, WT- or N153Q-infected IFNAR^-/-^ mice collected at day 5 pi. (B) Kaplan-Meier survival curve. (C) Body weight loss profile. (D) Clinical scores. (E) Viremia titers were measured by plaque assay at day 4 and 6 pi. (n=3-4). Dotted line indicates the detection limit of the assay. Data are representative of at least two independent experiments. Statistical analysis was performed using Mann–Whitney U test. *, p < 0.05; ns, not significant.

Heterologous passive transfer experiments were also conducted where N153Q serum was administered to WT-infected muMT IFNAR^-/-^ mice, and vice versa. Transfer of N153Q and WT serum did not confer significant protection to WT- and N153Q mutant-infected muMT IFNAR^-/-^ mice, respectively, as evidenced by 100% mortality, sharp body weight loss profile and severe clinical manifestations (Fig. 4B-D). This was accompanied by moderate and transient reduction in viremia titers at day 4 pi. but not at day 6 pi. (Fig. 4E). These observations thus implied that N153Q mutant was susceptible to clearance by antibodies only present in N153Q serum.

To identify the antibody class that was involved in N153Q mutant accelerated clearance in IFNAR^- /-^ mice, we monitored the DENV-specific IgM and IgG titers until day 6 pi. Domain III on E protein (EDIII) was used as coating antigen in the ELISA as it is unaffected by the N153Q mutation which is located on EDI. Comparable anti-EDIII IgM antibody titers were detected in sera from WT- and N153Q mutant-infected IFNAR^-/-^ mice from day 4 pi. onwards, which were significantly higher than the baseline titers measured in non-infected serum samples (Fig. 5A). Expectedly for such early phase of the infection, the anti-EDIII IgG levels detected in both infected groups were not significantly different from the levels measured in uninfected serum samples (Fig. 5B).

**Figure 5.**
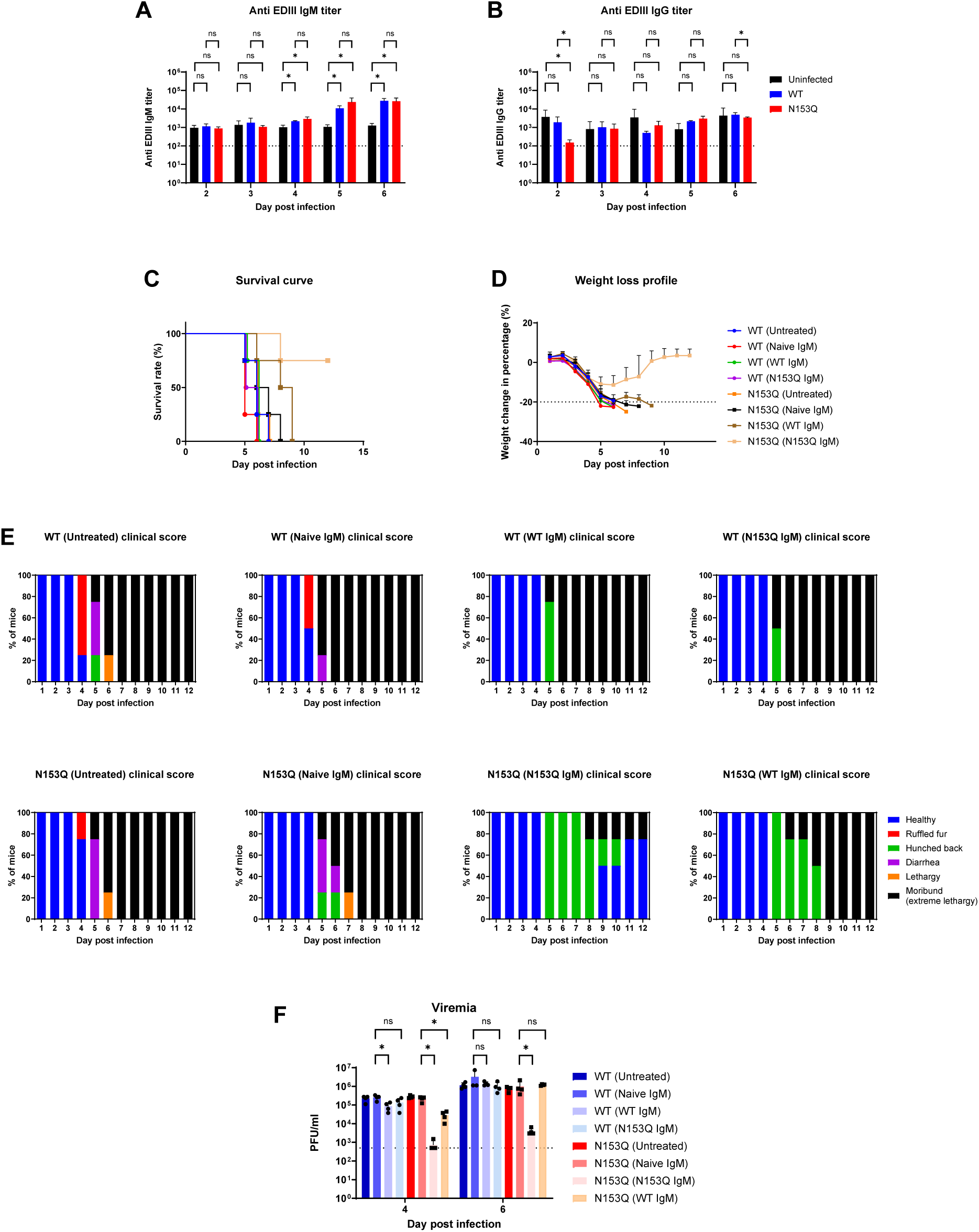
Homologous and heterologous passive transfer of purified IgM in muMT IFNAR^-/-^ mice. 5-7 weeks old IFNAR^-/-^ mice (n=4) were sc. infected with WT or N153Q DENV2 at 10^5^ PFU/mouse. Serum anti-DENV EDIII IgM (A) and (B) IgG titers were monitored by ELISA, where purified EDIII protein was used as coating antigen. (C-F) 5-7 weeks old muMT IFNAR^-/-^ mice (n=4) were sc. infected with WT or N153Q DENV2 at 10^5^ PFU/mouse. At day 3 pi. mice were administered intravenously with 30ug of total IgM purified from naïve, WT- or N153Q-infected IFNAR^-/-^ mice at day 5 pi. (C) Kaplan-Meier survival curve. (D) Body weight loss profile. (E) Clinical scores. (F) Viremia titers were measured by plaque assay at day 4 and 6 pi. Dotted line indicates the detection limit of the assay. Data are representative of at least two independent experiments. Statistical analysis was performed using Mann–Whitney U test. *, p < 0.05; ns, not significant.

We next purified total IgM antibodies from serum samples harvested at day 5 pi. of WT- and N153Q mutant-infected IFNAR^-/-^ mice and passively administered equal amounts to WT- and N153Q mutant-infected muMT IFNAR^-/-^ mice, in homologous and heterologous combinations. The results obtained mirrored the outcome of passive transfers performed using serum samples, whereby only homologous passive transfer of IgM purified from N153Q mutant-infected IFNAR^- /-^ mice protected N153Q mutant-infected muMT IFNAR^-/-^ mice and reduced the viremia titers; while WT virus remained resistant to WT IgM- and N153Q IgM-mediated clearance, and N153Q virus was resistant to WT IgM-mediated clearance (Fig. 5C-F). Hence, these results strongly suggested that DENV N153Q-specific IgM were responsible for the accelerated clearance of N153Q mutant in IFNAR^-/-^ mice.

To gain further mechanistic insights, we investigated whether N153Q mutant was more susceptible to IgM-mediated neutralizing activity. PRNT_50_ assays were conducted using WT and N153Q viruses produced in either mosquito or mammalian cell lines. Results indicated that day 5 pi. serum from WT and N153Q-infected IFNAR^-/-^ mice displayed significant and comparable neutralizing activities against WT and N153Q viruses, respectively (Suppl. Fig. S6), hence ruling out that N153Q mutant may be more susceptible to IgM-mediated neutralization than WT virus. Notably, the neutralizing activities were much reduced when viruses were produced in mammalian cells, supporting that glycan complexity influences antibody-mediated neutralization.

Altogether, these results indicated that N153Q mutant’s attenuated phenotype in IFNAR^-/-^ mice characterized by accelerated clearance in the blood circulation was due to N153Q-specific non-neutralizing IgM antibodies.

### N153-linked glycans interfere with IgG-mediated neutralization

While the data generated in the above sections strongly supported that N153-linked glycans on E protein protected DENV from premature clearance by non-neutralizing IgM antibodies early during infection, we next explored the potential role of these glycans at a later stage of the infection, when IgG antibodies are produced, or during secondary homologous infection. However, as WT-infected IFNAR^-/-^ mice had to be euthanized by day 7 pi., we turned to human plasma samples from convalescent dengue patients.

Both WT and N153Q viruses produced in mosquito cells were equally neutralized by DENV2 convalescent plasma samples (Fig. 6A, left panel), supporting that N153Q mutant was not more susceptible to IgG-mediated neutralization than WT DENV, and implying that N153-linked glycans did not influence IgG-mediated neutralization of DENV. However, when PRNT_50_ assays were conducted with viruses produced in mammalian cells, N153Q mutant was more effectively neutralized than WT virus, and this was particularly evident in Vero cells (Fig. 6A, middle panel), less so in Huh-7 cells (Fig 6A, right panel). Furthermore, as observed with day 5 pi. mouse serum samples (Suppl. Fig. S6), the overall neutralizing activity of convalescent plasma samples against mosquito cell-produced DENV was significantly higher than that seen with mammalian cell-produced DENV (Fig. 6A). Similar observations were made with plasma samples from DENV1 convalescent patients that neutralized more effectively N153Q DENV1 mutant than its WT counterpart, although the differences were not statistically significant (Fig. 6B). Together, these observations indicated that the role of N153-linked glycans in shielding DENV from IgG-mediated neutralization was dependent on the cell line used to produce the virus and suggested that glycan complexity on E protein at both N67 and N153 sites influenced IgG-mediated neutralizing activity.

**Figure 6.**
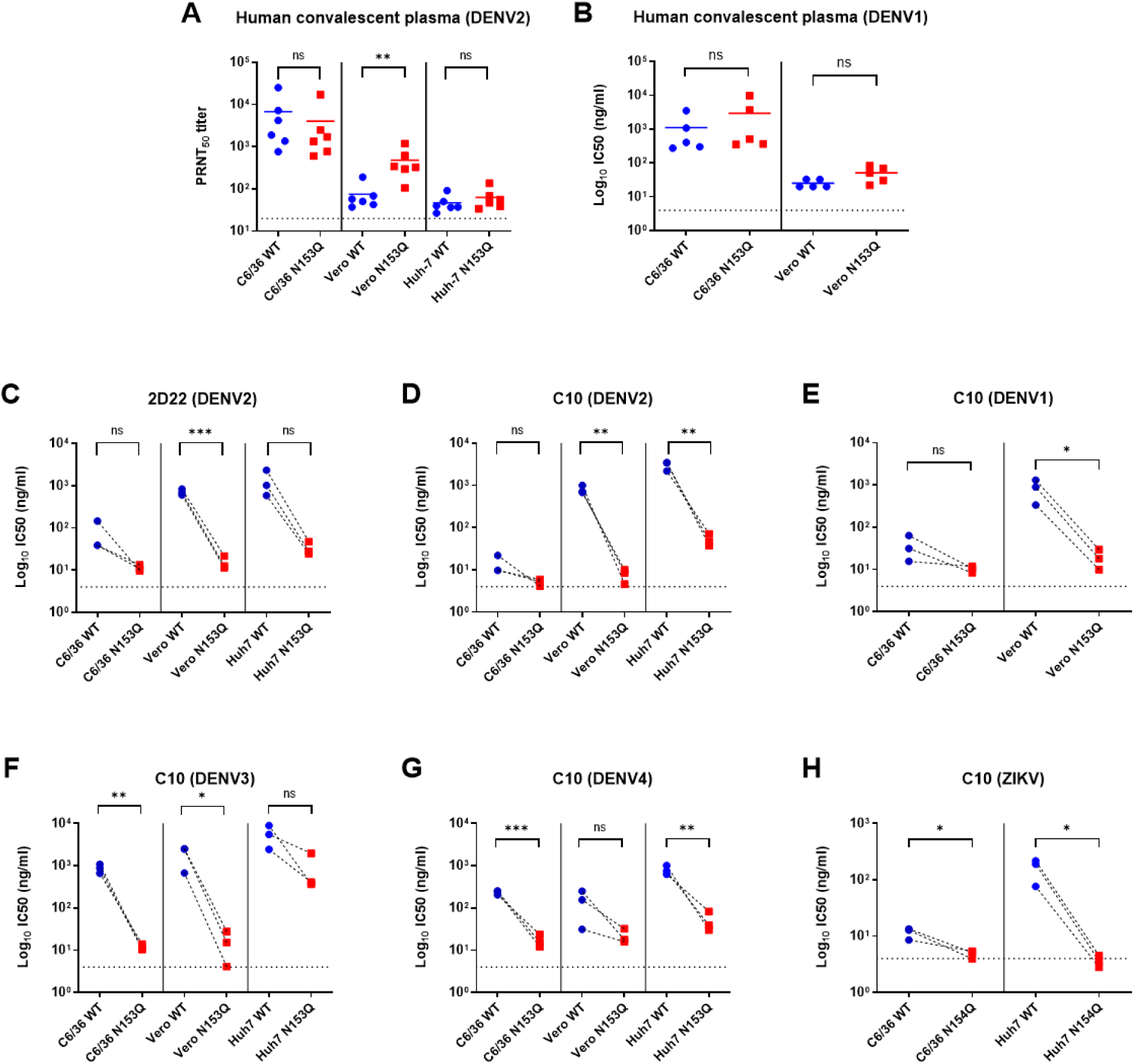
Sensitivity of N153Q DENV1-4 and N154Q ZIKV mutants to neutralization by convalescent plasma and monoclonal antibodies. Neutralizing activity of (A) DENV2 (n=6) and (B) DENV1 (n=5) human convalescent plasma against corresponding WT and N153Q DENV produced in C6/36, Vero or Huh-7. (C) IC_50_ values of 2D22 mAb against WT DENV2 and N153Q DENV2 mutant. (D-H) IC_50_ values of C10 mAb against (D) WT DENV2 and N153Q DENV2 mutant; (E) WT DENV1 and N153Q DENV1 mutant; (F) WT DENV3 and N153Q DENV3 mutant; (G) WT DENV4 and N153Q DENV4 mutant; (H) WT ZIKV and N154Q ZIKV mutant. For C to H, each data point represents mean values of an independent experiment. Dotted line indicates the detection limit of the assay. Statistical analysis was performed using Mann–Whitney U test (A) and t-test (B&C). *, p< 0.05; **, p < 0.01; ns, not significant.

To further investigate whether N153-linked glycans on E protein of DENV influence IgG-mediated neutralization, we evaluated the neutralizing activity of monoclonal antibodies (mAb) C10 and 2D22 against WT and N153Q DENV2 mutant. Both mAbs were previously reported to recognize two distinct conformational epitopes encompassing the N153-linked glycan loop, and induced disorder or change in position of the glycan loop on DENV and ZIKV, hinting that N153-linked glycans may interfere with the mAbs neutralizing activity [32–34, 45]. Consistently, the PRNT results indicated that absence of N153-linked glycans on E protein generally increased the susceptibility of DENV2 to mAb-mediated neutralization, as evidenced by the significant reduction in IC_50_ concentrations, and regardless of the cell type used for virus production (Fig. 6C-D). The enhanced susceptibility of N153Q mutant to neutralization by C10 and 2D22, compared to WT, was however less striking when the viruses were produced in mosquito cells compared to mammalian cell lines (Fig. 6C-D), in line with observations made with convalescent plasma samples. Specifically, the lack of N153-linked glycans on DENV2 increased the potency of C10 mAb by 3-fold, 105-fold, and 58-fold against the virus produced in C6/36, Vero, and Huh-7 cells, respectively; while the potency of 2D22 mAb increased by 7-fold, 47-fold, and 40-fold (Fig. 6C-D).

Similar trends were observed with the three other DENV serotypes and ZIKV, whereby the lack of N153/N154-linked glycans increased virus susceptibility to C10 mAb neutralization (Fig. 6E-H), implying that the findings can be extended to all DENV serotypes and other orthoflaviviruses. Overall, the lack of N153-linked glycans on E protein enhanced DENV and ZIKV susceptibility to mAb-mediated neutralization and was highly dependent on host cells used for virus propagation. These findings thus implied that glycoforms produced in different cell lines might influence antibody access and binding to conformational epitopes on E protein.

### Glycan bulkiness and complexity influence IgG binding to DENV

The neutralization assays described above suggested that glycan composition on DENV particles produced in different cell lines influenced IgG-mediated neutralization. To test this hypothesis, glycoproteomics was employed to determine the glycoforms at N67 and N153 on E protein purified from WT DENV2 propagated in C6/36, Vero and Huh-7. The results showed that N153-linked glycosylation site exhibited greater glycan species diversity than N67, regardless of the cell line used for virus production (Suppl. Fig. S6). Specifically, 14, 55, and 21 glycan species were detected at the N153-linked site compared to just 6, 7, and 9 glycan species at the N67-linked site when WT virus was produced in C6/36, Vero, and Huh-7 cells, respectively. The various glycoforms detected were then grouped into high-mannose, hybrid/complex, fucosylated, and sialylated glycans. This approach revealed distinct glycan composition patterns at each N-glycosylation site and depending on the cell line used for virus production. High-mannose glycans predominated at both N67- and N153-linked sites when the virus was grown in C6/36 cells, while lower levels were observed in Vero cells and Huh-7 cells (Fig. 7A-B). Conversely, hybrid/complex glycans were more abundant in Vero- and Huh-7-derived viruses, with the highest levels in Huh-7 cells (Fig. 7A-B). Sialylation was exclusively detected on both N-linked sites in viruses grown in Huh-7 cells (Fig. 7A-B). Notably, N153-linked glycans were highly fucosylated (50-70%), while fucosylated glycans were rare at the N67-linked site, regardless of the cell line used to produce the virus (Fig. 7A-B).

**Figure 7.**
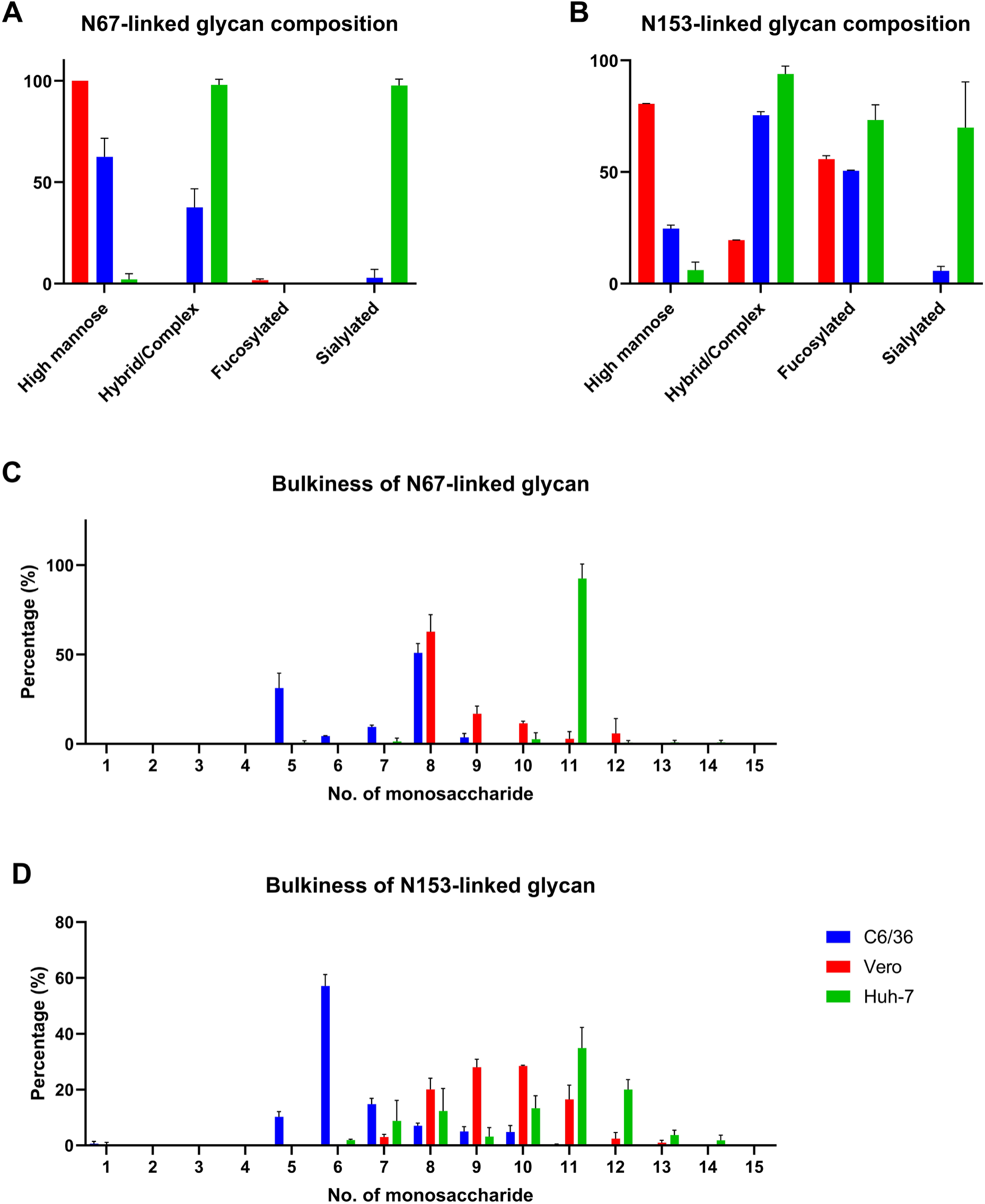
Composition of N-linked glycans at N67 and N153 of E protein from WT DENV2. (A) N67-linked and (B) N153-linked glycans on E protein of WT DENV2 produced in C6/36, Vero or Huh-7 cell lines were classified into high mannose, hybrid/complex, fucosylated and sialylated. The percentage of each glycan class was plotted. (C) N67-linked and (D) N153-linked glycans were grouped by bulkiness/number of monosaccharides on each glycoform. For example, HexNAc(2)Hex(3)Fuc(1) is counted as 6 monosaccharides. The percentage of glycan based on the number of monosaccharides was plotted. Data represent mean values of two independent biological replicates.

We also categorized the glycoforms based on their “bulkiness,” determined by the number of monosaccharide units per glycoform. The analysis revealed that most N67-linked glycans contained 5 or 8 monosaccharide units when the virus was produced in C6/36 cells (Fig. 8C). For Vero cell-produced virus, N67-linked glycans were primarily composed of 8 monosaccharides, although a significant proportion had 9-12 units (Fig. 8C). N67-linked glycans from virus produced in Huh-7 cells contained 11 monosaccharides, making them the bulkiest across the cell lines (Fig. 8C). This pattern suggested that glycan bulkiness at N67 position increased progressively from C6/36 to Vero to Huh-7 cells. A similar trend was noted for N153-linked glycans, with the majority containing 6 monosaccharides in C6/36 cells, increasing to 8-11 in Vero cells and 10-12 in Huh-7 cells (Fig. 8D).

**Figure 8.**
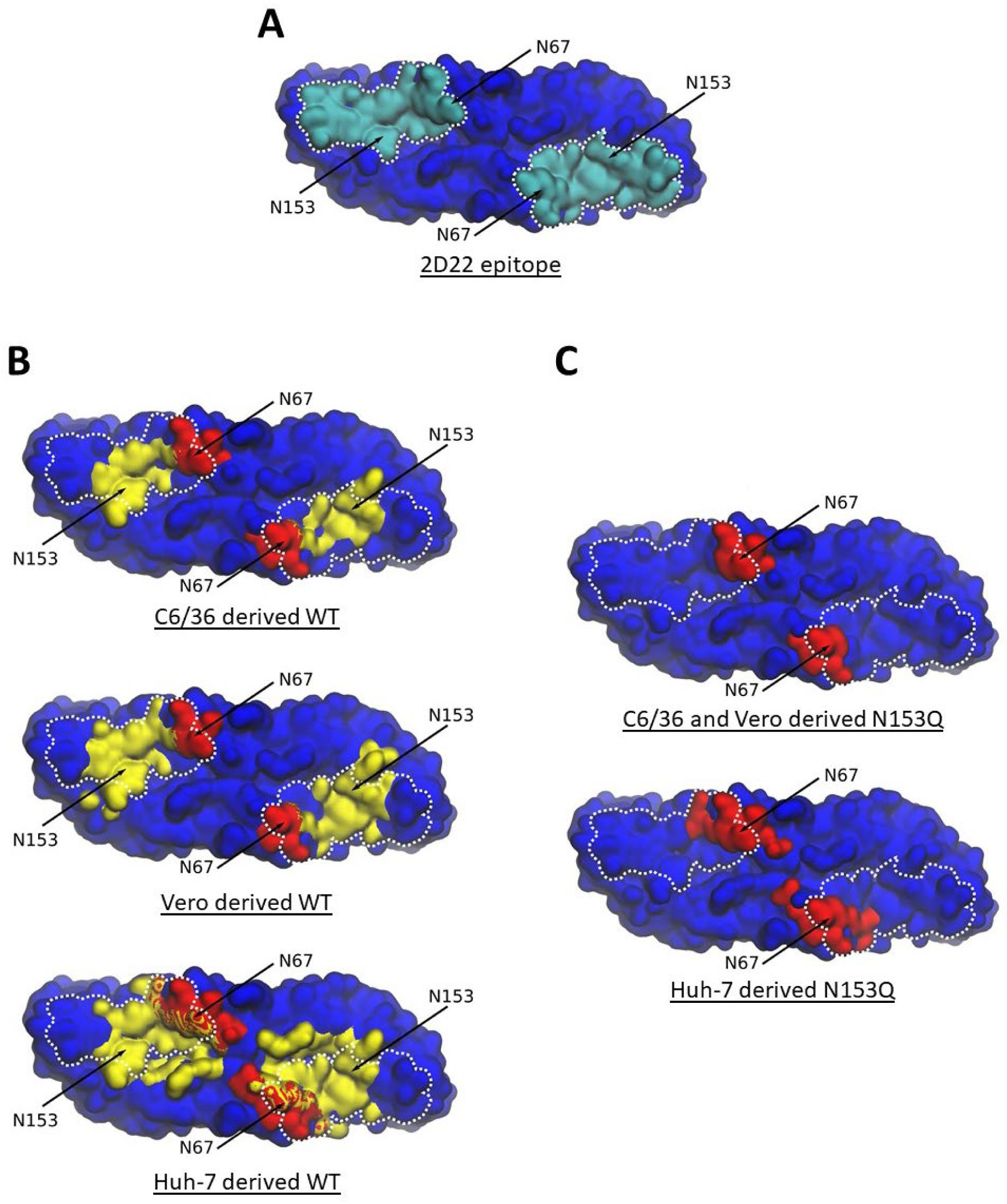
Molecular dynamics simulation of N-linked glycans shielding effect. (A) mAb 2D22 epitope surface was colored in cyan and delineated by white dots. (B) Amino acid residues of E protein ectodomain interacting with N-linked glycans on WT DENV2 produced in C6/36, Vero or Huh-7 cells. (C) Amino acid residues of E protein ectodomain interacting with N67-linked glycans on N153Q DENV2 produced in C6/36 or Huh-7 cells. As the N67-linked glycan composition was identical on viruses grown in C6/36 and Vero, only simulation in C6/36 was performed. Amino acid residues interacting with N67-linked and N153-linked glycans were colored in red and yellow, respectively while amino acid residues interacting with both N-linked glycans were colored in striped red and yellow.

All atom explicitly solvated molecular dynamics (MD) simulations were conducted to explore the spatiotemporal dynamics of glycans shielding the epitope on E protein dimer which can interfere with IgG-mediated neutralization. For the simulations, the predominant glycoforms detected by glycoproteomics at each N-glycosylation site for all cell lines were used (Suppl. Fig. S7). The 2D22 mAb footprint on the ectodomain of E protein was mapped using previously published cryoEM data [32] and was found to encompass both N67 and N153 residues (Fig. 8A). Our simulations showed that glycans at both N67 and N153 sites interacted with amino acid residues that significantly overlapped with 2D22 binding footprint, across all the cell lines (Fig. 8B). However, the absence of N153-linked glycans in the case of N153Q mutant exposed a substantial surface area for mAb 2D22 binding (Fig. 8C). Simulations indicated that the bulkier the glycans at either N-sites on mammalian cell line-produced virus (Vero and Huh-7) the greater the surface of interactions with E protein ectodomain compared to glycans derived from mosquito cells (C6/36), suggesting greater interference with the antibody binding (Fig. 8B&C). The specific amino acid residues interacting with N-linked glycans were identified, reinforcing the idea that bulkier glycans shield more of the E protein surface (Suppl. Table S3).

Our *in-silico* study supports that glycans at both N67 and N153 sites on E protein interfered with 2D22 mAb binding to its conformational epitope, with bulkier glycans interacting with a larger surface area, thus contributing to greater steric hindrance and reduced Fab binding. These findings help explain why N153Q mutant produced in mosquito cells was more susceptible to 2D22 neutralization than N153Q produced in mammalian cells (Fig. 6C).

## Discussion

Glycan structures have been shown to play an important role in the pathogenesis of many viruses, by influencing virus receptor binding, inducing immune responses and masking antigenic sites. The envelope glycoprotein gp120 of human immunodeficiency virus (HIV) for example is heavily glycosylated (more than 90 sites per monomer), forming a glycan shield that masks critical epitopes from being recognized by neutralizing antibodies [46, 47]. This glycan shield on HIV particles has been found to be very dynamic with rapid deletion and addition of N-linked glycosylation sites, making difficult the design of Envelope-based vaccine candidates able to protect against all HIV glycovariants produced over time within a single infected individual. Similarly, the hemagglutinin (HA) protein of the influenza virus is heavily glycosylated and undergoes rapid changes in glycosylation patterns and sites, which can alter antigenic sites, contributing to antigenic drift, and allowing the virus to evade immune detection [48].

In sharp contrast, DENV Envelope protein contains only two N-linked glycosylation sites (N67 and N153) that are highly conserved across the four serotypes of DENV. The highly conserved nature of these two N-linked glycosylation sites implies a strong selective pressure and a critical role in DENV fitness and virulence. Here, we demonstrated the role of N153-linked glycans in protecting the virus from antibody-mediated clearance, which represents a novel immune evasion mechanism for DENV. Uniquely, we showed that during the early stage of infection, N153-linked glycans prevented premature viral clearance by non-neutralizing IgM antibodies. We hypothesize that the lack of N153-linked glycans could increase IgM antibody binding efficacy at the virus surface, thereby enhancing Fc receptor-dependent phagocytosis of virus-antibody complexes. IgM antibodies interact with Fc receptors such as pIgR, FcμR, and Fcα/μR [49, 50]. Given its limited role in DENV pathogenesis, pIgR likely does not contribute to the observed attenuation [49]. FcμR, primarily expressed on murine B cells, could potentially influence viral clearance; however, our data from muMT IFNAR^-/-^ mice lacking mature B cells suggest that FcμR-bearing B cells are not involved, while the involvement of FcμR-bearing non-B cells would require further investigation [51–53]. The role of Fcα/μR, found on macrophages, dendritic cells, and other hematopoietic cells, remains a possibility, as Fcα/μR has been implicated in IgM-mediated pathogen clearance through endocytosis [49, 54]. The role of FcμR-bearing non-B cells or Fcα/μR on macrophages and dendritic cells remains unexplored and may warrant further study to determine their influence on IgM-mediated clearance of N153Q mutant.

Alternatively, the N153Q mutant may be more susceptible to IgM-initiated complement activation through the classical pathway, culminating to opsonin (C3b and C5b) deposition and membrane attack complex (MAC)-mediated virolysis. One could speculate that N153-linked glycans on DENV E protein shield the virus from opsonin deposition and MAC formation on DENV phospholipid bilayer. Complement-mediated virolysis has indeed been reported in other enveloped viruses. In West Nile virus (WNV), also a member of the orthoflavivirus genus, C3b deposition was reported to inhibit viral fusion with the host cell plasma membrane [22]. C3 deposition was also observed on the surface of a paramyxovirus, Simian Virus 5, which led to extensive aggregate formation [55, 56]. Furthermore, MAC-mediated virolysis has been reported to effectively neutralize HIV [57, 58]. However, direct evidence of the role of N-linked glycans on DENV or other viruses in interfering in complement-mediated virolysis is currently lacking and warrants further investigation.

In addition to preventing IgM-mediated viral clearance, we also showed that N153-linked glycans on E protein of DENV antagonized IgG-mediated neutralization. Previous structural studies suggested that these glycans might indeed interfere with mAbs 2D22 and C10 binding [32–34, 45]. 2D22, a DENV2 specific mAb, binds both the EDIII and glycan loop (residues 140-160) containing N153-glycosylation site on DI of one E subunit, and the fusion loop on DII of another subunit. As such, 2D22 was shown to inhibit fusion by preventing E protein dimer to trimer transition [32]. C10, a pan-orthoflavivirus broadly neutralizing mAb, also targets multiple domains of E protein and induces conformational change of the N153 glycan loop [33, 34, 45]. Combining biological (neutralizing assays), biochemical (glycoproteomics), and *in-silico* (MD simulation) approaches, we provided here direct and strong experimental evidence that N153-linked glycans shield DENV1-4 and ZIKV from IgG binding and neutralizing activity.

Our data also suggested that glycoforms on E protein influenced DENV resistance to antibody-mediated neutralization. DENV grown in mammalian cells was less susceptible to human plasma and mAb neutralization, and we proposed that bulkier, hybrid/complex glycans in mammalian-derived viruses enhanced resistance to neutralization. The relationship between glycan composition and/or glycan bulkiness and its biological functions such as shielding from neutralizing antibody has never been reported before, neither for DENV nor for other viruses. Although this shielding hypothesis is supported by *in-silico* modeling, experimental validation of the relationship between glycan bulkiness and immune evasion is needed to confirm its biological relevance.

Besides its main role in shielding the viral surface from antibody-mediated biological activities, we also observed that N153-linked glycans influenced organ tropism, as evidenced by lower viral titers specifically in the kidneys and liver from N153Q mutant-infected mice. This observation may be explained by the reduced infectivity of N153Q mutant in DC-SIGN, L-SIGN or mannose receptor bearing endothelial cells in these organs [37, 59–61], and correlated with the slight attenuation of this virus in BHK-21 and Huh-7 mammalian cell lines.

In conclusion, our study has uncovered a novel immune evasion strategy deployed by DENV and ZIKV to evade antibody-mediated clearance. In combination with other attenuating genetic alterations, the N153Q mutation represents a potential strategy for developing next generation live-attenuated DENV/ZIKV vaccines. Additionally, our findings support that host glycosylation pathways may be targeted therapeutically to alter the virus glycan composition, enhancing its susceptibility to neutralizing antibodies. Together, these novel insights into the role of N153/154-linked glycans not only deepen our understanding of DENV/ZIKV pathogenesis but also open avenues for innovative vaccine and therapeutic strategies.

## MATERIALS AND METHODS

### Ethics statement

All the animal breeding and experimental procedures were approved by the National University of Singapore (NUS) Institutional Animal Care and Use Committee (IACUC) under protocol numbers BR16-0156, BR22-0139, R16-0422 and R18-1400. Animal facilities are licensed by the regulatory body Agri-Food and Veterinary Authority of Singapore. The experiments involving human samples were reviewed and approved by Domain Specific Review Board (DSRB) under reference no. 2008/00293 (DENV2) and E/2009/432 (DENV1) [62].

### Cell lines and culture condition

Baby hamster kidney fibroblasts BHK-21 (ATCC # CCL-10) were cultured in Roswell Park Memorial Institute (RPMI) 1640 medium (GIBCO, cat. # 22400) with 10% fetal bovine serum (FBS) (GIBCO, cat. # 10270106) at 37°C with 5% CO_2_ in a humidified incubator. African green monkey kidney epithelial cells Vero (ATCC # CCL-81™) and human hepatocellular carcinoma Huh-7 cells (a kind gift from Prof Ooi Eng Eong, Duke-NUS, Singapore) were cultured in 10% FBS Dulbecco’s Modified Eagle Medium (DMEM) (GIBCO, cat. # 11965) at 37°C with 5% CO_2_ in a humidified incubator. *Aedes albopictus* larvae cells C6/36 (ATCC # CRL-1660™) were maintained in Leibovitz’s L-15 medium (GIBCO, cat. # 11415) with 10% FBS at 28°C in closed-cap flasks. All cell lines were routinely tested for mycoplasma contamination using a detection kit (InvivoGen, cat. # rep-mys).

### Viruses and propagation condition

DENV1 05K3903DK1 (GenBank accession number: EU081242.1) is a clinical isolate obtained during the 2005 dengue outbreak in Singapore [63]. DENV2 D2Y98P (GenBank accession number: JF327392) was isolated in Singapore in 1998 and plaque purified in BHK-21 cells, followed by generation of an infectious clone [64]. DENV3 C0360/94 (GenBank accession number: KJ737429.1, cat. # NR-48800) and DENV4 703-4 (cat. # NR-48801) were both isolated in Thailand in 1994 and were sourced from the Biodefense and Emerging Infections Research Resources (BEI Resources) repository. ZIKV H/PF/2013 (GenBank accession number: KJ776791.2) was isolated in French Polynesia in 2013.

All viruses were propagated by infecting cell monolayers (80-90% confluent) in tissue culture flasks, followed by a 1-hour incubation with gentle rocking at 15-minute intervals to facilitate viral adsorption. Afterward, the appropriate medium was added, and the infected flasks were incubated for several days. For propagation in C6/36 cells, the infected cells were maintained in 2% FBS Leibovitz’s L-15 medium in closed-cap tissue culture flasks at 28°C. For Vero and Huh-7 cells, the infected cultures were maintained in 2% FBS DMEM at 37°C with 5% CO_2_ in a humidified incubator. Once 70-80% of the infected cells exhibited cytopathic effects (CPE), the culture supernatant was collected, centrifuged at 12,000 x *g* to remove cell debris, and stored at -80°C. Virus titers were determined by plaque assay.

### Generation of deglycosylated E protein DENV and ZIKV mutants

N153Q DENV2 mutant was previously described by us [65]. To generate N153Q DENV1,3,4 and N154Q ZIKV mutants, viral RNA from was first extracted using the QIAamp Viral RNA Extraction Kit (Qiagen, cat. # 52906) and reverse transcribed to cDNA using the GoScript Reverse Transcriptase Kit (Promega, cat. # A5000). The viral genome fragments were amplified by PCR with Q5® Hot Start High-Fidelity DNA Polymerase (New England Biolabs, cat. # M0494L). Primers used for cDNA synthesis and PCR were listed in Tables 1-5. PCR products were analyzed by gel electrophoresis, extracted using the QIAquick Gel Extraction Kit (Qiagen, cat. # 28704), and cloned into the pCR™Blunt II-TOPO™ vector, then transformed into One Shot™ TOP10 Chemically Competent E. coli (Invitrogen, cat. # K280020). Plasmids were verified by sequencing (Axil Scientific) using universal primers.

These plasmids that contain DENV1, DENV3, DENV4, and ZIKV genome fragments were used as templates for PCR. The N153Q/N154Q mutation was introduced into the E gene. The vector pACYC177-CMV-SV40pA-HDVR (kind gift from Prof. Ooi Eng Eong) was amplified and linearized by PCR. PCR products were gel purified as described, and viral constructs were assembled via Gibson assembly using NEBuilder® HiFi DNA Assembly Master Mix (New England Biolabs, cat. # E2621). In brief, equal molar ratio (0.2 pmol) of each fragment and vector which are flanked with overlapping sequences, were seamlessly assembled at 50°C for 2 hours. The final plasmid contained the CMV promoter, viral genome, HDV ribozyme, and SV40 poly-A sequence.

BHK-21 cells were seeded at 2.5 × 10⁵ cells/well in a 6-well plate with 10% FBS Minimum Essential Medium (MEM) (GIBCO, cat. # 11095) a day before transfection. Cells were washed and replenished with Opti-MEM (Gibco, cat. # 31985062). The assembled viral constructs were transfected using Lipofectamine™ 2000 Transfection Reagent (Invitrogen, cat. # 11668019). After 4 hours, the medium was replaced, and supernatants from days 4-6 post-transfection were collected, centrifuged at 12,000 × g for 10 mins at 4°C, and stored at -80°C. Viruses were propagated in C6/36 cells and titrated by plaque assay. RNA was extracted, converted to cDNA using specific primers, amplified by PCR, and sequenced.

### In vitro viral infection kinetic assay

C6/36 cells were seeded in 96-well plates at 3.4 × 10⁴ cells per well, while BHK-21, Vero, and Huh-7 cells were seeded at 1.7 × 10⁴ cells per well. The cells were cultured in their respective media, supplemented with 10% FBS, and incubated overnight. The next day, cells were infected with viruses at a multiplicity of infection (MOI) of 0.1. Plates were incubated at 28°C (C6/36 cells), or at 37°C (Huh-7 and BHK-21 cells), for 1 hour, with rocking every 15 minutes to facilitate viral adsorption. Afterward, cells were gently washed twice with PBS to remove any unbound virus, and 200 µL of the respective medium supplemented with 2% FBS was added. Culture supernatants were collected at the specified time points and stored at -80°C. Four technical replicates were performed for each time point, and viral titers were determined using a plaque assay.

### Mouse models

IFNAR^-/-^ mice (C57BL/6 deficient in Type I interferon receptor) were obtained from The Jackson Laboratory (strain # 010830). muMT mice (C57BL/6 deficient in mature B cells, strain # 002288) were generously provided by Prof. Laurent Renia, Lee Kong Chian School of Medicine, NTU. To generate muMT IFNAR^-/-^ mice, muMT/muMT mice were crossed with IFNAR^-/-^ mice to produce heterozygous muMT/+ IFNAR^+/-^ offspring. These were then inbred to obtain muMT/muMT IFNAR^-/-^ mice (referred to as muMT IFNAR^-/-^). Genotyping was carried out to identify mice with the desired genotype using quick DNA purification and genotyping protocols from The Jackson Laboratory. Briefly, a 1-2mm segment of the mouse tail was cut and incubated in alkaline lysis buffer (25mM NaOH and 200nM disodium EDTA). Samples were heated to 95°C for 1 hour, then neutralized with neutralizing buffer (40mM Tris). PCR amplification was done using DreamTaq™ Hot Start Green PCR Master Mix (Thermo Fisher Scientific, cat. # K9021), following the thermocycling conditions specified by The Jackson Laboratory.

### Mouse infection

Five to seven weeks old mice were subcutaneously (sc.) infected with 10^5^ or 10^6^ PFU/mouse of virus propagated in C6/36 as indicated in the figure legend. SC injection was performed under loose skin around the neck and shoulder area of the mouse. The infected mice were monitored daily for weight loss and clinical scores or were bled for viremia quantification. Mice were euthanized when body weight loss reached 20% (DENV2) or 30% (ZIKV) of initial weight (at day 0) or when they were moribund, whichever came first.

### Serum and organ collection

Blood samples were collected at the specified time points from the retro-orbital sinus using a microhematocrit heparinized capillary tube (Fisher Scientific, cat. # 22-362566) under isoflurane anesthesia. The blood was then centrifuged at 6,000 × *g*, 10°C for 10 minutes. Serum was separated and stored at -80°C. Viremia was quantified using a plaque assay. Mice were euthanized with a CO_2_ overdose and subsequently perfused with PBS to remove any residual blood from the organs. The organs were harvested and snap-frozen in liquid nitrogen, then stored at -80°C. The wet weight of each organ was recorded before they were homogenized using a tissue homogenizer in FBS-free RPMI-1640. The homogenates were clarified by centrifugation at 10,000 × g, and the supernatant was filtered through a 0.22 µm syringe filter. The filtered supernatant was serially diluted (5-fold) and viral titers were determined via plaque assay. Viral loads in the organs were normalized to the wet weight of the tissues and expressed as PFU/g.

### B cell depletion

Five to seven weeks old IFNAR^-/-^ mice were intraperitoneally (ip.) administered with 100ug/mouse of anti-mouse CD20 antibody (Bio X Cell, cat. # BP0356) or mouse IgG isotype control (Bio X Cell, cat. # BE0093) 2 days prior to DENV infection.

### IgM purification and passive transfer

Five to seven-week-old IFNAR^-/-^ mice were sc. infected with 10⁵ PFU/mouse. Serum was collected from the infected IFNAR^-/-^ mice on day 5 pi., and then heat-inactivated at 56°C for 30 minutes. IgM antibodies were purified from the heat-inactivated sera using a commercial IgM purification kit (Advanced BioReagents Systems, cat. # C8056-M). The total IgM concentration was measured using an ELISA kit (Abcam, cat. # ab133047). Five to seven-week-old muMT IFNAR^-/-^ mice were sc. infected with 10⁵ PFU/mouse and, on day 3 pi., were intravenously (iv.) administered with 200µL of neat serum or 30ug purified IgM per mouse.

### Blood parameters measurement

Mouse blood samples were collected in MiniCollect™ tubes containing K2EDTA (Greiner, cat # 450480). The whole blood was promptly analyzed for cell counts using the automated Cell-Dyn 3700 Hematology Analyzer (Abbott) by NUS Comparative Medicine. This analyzer employs fluorescent flow cytometry and advanced analytical algorithms for accurate blood cell counting.

### Inflammatory cytokines measurement

Mouse blood was collected in MiniCollect™ tubes containing the anticoagulant K2EDTA and centrifuged at 6,000 × *g*, 4°C for 10 minutes. The plasma was then collected for cytokine measurement using the LEGENDplex Mouse Inflammatory Panel kit (BioLegend, cat. # 740446), following the manufacturer’s instructions. Samples were analyzed using an Attune Nxt flow cytometer (Thermo Fisher Scientific) and data were processed with FlowJo software.

### Bulk RNA-seq of mouse white blood cells

To isolate white blood cells, mouse blood was collected in MiniCollect™ tubes containing the anticoagulant K2EDTA and centrifuged at 6,000 × *g*, 4°C for 10 minutes. The plasma was removed, and the blood cells were resuspended in red blood cell lysis buffer (4.13g NH4Cl, 0.5g KHCO3, and 100µL of 0.5M EDTA in 500mL of water) at a 1:10 ratio of whole blood to lysis buffer. The mixture was incubated at room temperature for 5 minutes, after which the lysis was stopped by adding 10% FBS RPMI-1640 at a 1:2 ratio of lysis buffer to RPMI-1640. The mixture was then centrifuged at 700 × *g* for 10 minutes at 4°C, and the supernatant was carefully discarded. The cell pellet was washed twice with 10% RPMI-1640.

To extract total RNA, the cell pellet was mixed with 1mL of TRIzol™ reagent (Invitrogen, cat. # 15596026) and incubated at room temperature for 5 minutes. Next, 200µL of chloroform was added, and the mixture was shaken vigorously for 15 seconds before incubating for 2–3 minutes. The solution was then centrifuged at 12,000 × *g* for 15 minutes at 4°C. The aqueous phase was transferred to a clean tube and mixed with 1mL of 100% isopropanol, followed by incubation at room temperature for 10 minutes. The solution was passed through RNeasy spin column (Qiagen, cat. # 74104) by centrifugation at 8,000 × *g* for 30 seconds. The spin column was washed sequentially with 700µL of RW1 buffer once and 500µL of RPE buffer twice. It was then dried by centrifugation at maximum speed for 1 minute. RNA was eluted with water.

RNA samples were sent to Azenta Life Sciences for bulk RNA sequencing. In brief, total RNA was subjected to polyA capture for mRNA enrichment during library preparation. Sequencing was performed on the Illumina NovaSeq platform in a 2×150bp paired-end configuration, generating approximately 15.0 Gb of data (50 million reads) per sample. Raw data analysis was conducted by Dr. Chan Kuan Rong from EID, Duke-NUS Medical School. Raw counts from the RNA-seq count matrices were imported into Partek Genomics Suite v7.21, where one-way ANOVA and t-tests were performed for comparisons between WT vs PBS, N153Q vs PBS, and N153Q vs WT. Differentially expressed genes (DEGs) were identified using the Python packages Pandas (v1.5.2) and NumPy (v1.23.5), based on Python 3.9.7. Data visualization and DEG analysis were facilitated using STAGEs, with log2 fold-change ≥ 1 and p-value < 0.05 as cutoff criteria [66]. Volcano and bar plots were generated using Prism 9.4.0.

### Enzyme-linked immunosorbent assay (ELISA)

Systemic DENV-specific IgM and IgG antibody titers were determined using an indirect ELISA. 96-well EIA plates (Corning, cat. # 3590) were coated overnight at 4°C with either polyethylene glycol (PEG)-purified WT DENV2 propagated in Vero cells (150ng/well) or purified WT DENV2 EDIII protein (250ng/well), diluted in PBS. The plates were washed three times with wash buffer (0.05% Tween-20 in PBS) and then incubated with blocking buffer (5% bovine serum albumin in wash buffer) for 1 hour at 37°C. After decanting the blocking buffer, serially diluted (2 to 5-fold) samples were added to the wells, and the plates were incubated for 1 hour at 37°C, followed by three washes with wash buffer. Secondary antibodies, diluted in blocking buffer, were then added and incubated for 1 hour at 37°C. Goat anti-mouse IgM conjugated to horseradish peroxidase (HRP) (Bio-Rad, cat. # 31440, 1:1,000 dilution) or goat anti-mouse IgG HRP (Bio-Rad, cat. # 170-6516, 1:3,000 dilution) were used as secondary antibodies. After decanting the secondary antibody, the plates were washed three times with wash buffer. 3,3’,5,5’-Tetramethylbenzidine (TMB) solution (Invitrogen, cat. # 00-4201-56) was added to the wells and incubated for 10 minutes at room temperature. The reaction was stopped by adding 1M H₂SO₄, and absorbance was measured at 450nm.

### Plaque reduction neutralizing (PRNT) assay

Plasma or serum samples were heat-inactivated at 56°C for 30 minutes and then serially diluted (2 to 5-fold) in a virus suspension containing approximately 500 PFU/mL, diluted in RPMI-1640 with 2% FBS. The antibody-virus mixtures were incubated at 37°C for 1 hour before being titrated by plaque assay. The percentage of neutralization was calculated by comparing the number of plaques in the antibody-treated samples to those in samples with only virus and medium. The PRNT_50_ titer was defined as the plasma/serum dilution that resulted in 50% plaque neutralization. Monoclonal antibodies 753[3], C10 (Creative Biolabs, cat. # MRO-1607CQ), and 2D22 (Creative Biolabs, cat. # PABL-065) were tested in a similar PRNT assay, but without the heat inactivation step. The half-maximal inhibitory concentration (IC50) was determined as the concentration of antibody that resulted in 50% plaque neutralization compared to the virus-only control.

### Glycoproteomics analysis

C6/36, Vero, and Huh-7 cells were cultured to 70-80% confluency. On the day of infection, the culture medium was removed, and the cells were incubated with WT DENV2 diluted in FBS-free medium for 1 hour to allow viral adsorption. The cells were incubated at 28°C or 37°C, with or without 5% CO2, depending on the cell line. Culture supernatants containing viruses were collected between days 3 and 5 post-infection (pi), when 70-80% of the infected cells showed cytopathic effect (CPE). The harvested supernatant was subjected to a series of centrifugations at 4°C: 100 × *g* for 20 minutes, 4,000 × *g* for 35 minutes, 10,000 × *g* for 35 minutes, and 100,000 × *g* for 90 minutes. After the final centrifugation, the supernatant was discarded, and the pellet was resuspended in 100 mM Tris-HCl, pH 7.4 for virus grown in C6/36 cells or FBS-free medium for virus grown in Vero and Huh-7 cells.

For virus grown in Vero and Huh-7 cells, co-immunoprecipitation was performed using Dynabeads Protein G (Thermo Fisher Scientific, cat. # 10003D) to increase the purity of the virus, following the manufacturer’s protocol. Briefly, a DENV-specific monoclonal antibody (Thermo Fisher Scientific, cat. # MA1-27093) diluted in PBS was first incubated with Dynabeads Protein G. The mixture was rotated for 20 minutes at room temperature, then placed on a magnet to separate the beads. The supernatant was discarded, and the antibody-conjugated beads were washed gently with PBS. The beads were then washed twice with conjugation buffer (20 mM sodium phosphate monobasic and 0.15 M sodium chloride in water, pH adjusted to 7-9), and resuspended in 5 mM bis(sulfosuccinimidyl)suberate (BS3) (Thermo Fisher Scientific, cat. # 21580) in conjugation buffer. The mixture was rotated for 30 minutes at room temperature to allow antibody-bead crosslinking, then quenched with quenching buffer (3.025g Tris-HCl in 25mL water) and incubated for another 30 minutes at room temperature. Afterward, the supernatant was removed, and the beads were washed three times with PBS. The beads were resuspended in FBS-free medium containing virus and incubated with rotation for 2 hours at 4°C. The supernatant was discarded, and the beads were washed three times with PBS. The beads were then resuspended in PBS and transferred to a clean tube. After the supernatant was discarded, the beads were resuspended in cold 1% sodium dodecyl sulfate (SDS) diluted in PBS. The mixture was rotated for 10 minutes at room temperature, and the eluate containing purified virions was collected. All samples were incubated at 56°C for 30 minutes to inactivate the virus.

The samples were treated using the single-pot, solid-phase-enhanced sample preparation (SP3) protocol by Hughes et al [67]. In brief, 30 µg of protein were denatured and reduced using 8 M urea/50 mM Ambic and 10 mM DTT at 60 °C for 30 min followed by alkylation with IDA in the dark at room temperature for 30 min. The alkylation reaction was subsequently quenched by the addition of 15 µL of 10 mM DTT and incubation at room temperature for 15 min. Sera-Mag carboxylated speedbeads (Cytiva, Marlborough, MA) were added to the samples in 1:10 (wt/wt) protein/beads ratio. Acetonitrile was added to the suspension to a final concentration of 70 % ACN (v/v) and the final mixture was then incubated at 24 °C with mixing at 1200 rpm for 10 min. After 10 min, the magnetic beads were immobilized using a magnetic stand and the supernatant was removed. The magnetic beads were rinsed with 80 % ACN and the beads were immobilized again to discard the washing. This was repeated for another two times before the beads were reconstituted in 50 mM Ambic. Sequencing grade trypsin/lysC (Promega, Madison, WI) was added in 1:20 (wt/wt) enzyme/protein ratio to the beads and incubated at 37 °C for 16 hours. After 16 hours, the beads were centrifuged for 1 min at 20,000 g, immobilized on the magnetic stand and the supernatant collected into a clean tube. The supernatant was dried down using a vacuum centrifuge and stored at – 20 °C until the next step of analysis.

30ug of the sample after SP3 treatment were reconstituted in 30 µL of LCMS grade water and 10 µL of Sera-Mag carboxylated speedbeads (cat. no. 45152105050250) were added to the sample. Acetonitrile was then added to the mixture to a final concentration of 80 % (v/v). The suspension was incubated at 24 °C with mixing at 1200 rpm for 15 min. After 15 min, the magnetic beads were immobilized using a magnetic stand and the supernatant was removed. The magnetic beads were rinsed thrice with 80 % ACN. LCMS grade water was added to the beads and incubated at 24°C with mixing at 1,200 rpm for 15 min to elute the bound glycopeptides. The eluted glycopeptides were collected in a clean tube and dried down. The dried sample was reconstituted in LCMS water and injected into the LCMS for analysis.

18 ug of the glycopeptide enriched sample were injected onto a Dionex Ultimate RSLC 3000 nano LC system (Thermo Fisher Scientific, Waltham, MA) coupled to an Orbitrap Fusion Tribrid mass spectrometry system (Thermo Fisher Scientific, Waltham, MA) with an Easy-Spray™ nano source. The samples were trapped on a C18 trap column (Acclaim PepMap100 C18, 5 µm, 300 µm x 5 mm, Thermo Fisher Scientific, Waltham, MA) for 2 min before separation on an Easy-Spray™ C18 nano column (75 µm x 150 mm, 2 µm, Thermo Fisher Scientific, Waltham, MA) with a 107 min gradient at a flowrate of 0.3 µL/min. Solvent A was 0.1 % formic acid in water and solvent B was 0.1 % formic acid in ACN.

Mass spectrometry data was acquired in the positive ion data dependent acquisition (DDA) mode. Full scan MS spectra data were acquired in the mass range of m/z 500 to 2000, at an ionization source voltage of 2.2 kV, using the Orbitrap detector with resolution set to 120 K, normalized AGC target at 100 %, maximum injection time of 50 ms. MS/MS data was acquired using a top speed mode of 5 s in the Orbitrap detector at 30 K resolution, with an isolation window of 1.6 m/z, stepped HCD collision energy of 20 %, 30 % and 45 %, normalized AGC target of 200 % and maximum injection time of 100 ms. The criteria for MS/MS to be triggered for the precursor ions were minimum intensity of 5e4, charge states of 2 to 6. Dynamic exclusion was set at a duration of 15 s within a mass tolerance of 15 ppm.

Data analysis was performed using Thermo Proteome Discoverer (v2.2.0.388, Thermo Fisher Scientific, Waltham, MA) integrated with Byonic™ (v3.9.6, Protein Metrics, Cupertino, CA) node. The raw files were first matched against Swiss-Prot proteome database with the targeted NS1 protein sequence included. Software parameters were set as followed: missed cleavages set to two, digestion enzyme set to trypsin (full), precursor mass tolerance at 15 ppm, fragment mass tolerance at 20 ppm, fragmentation type to HCD, with modifications of methionine oxidations and deamidation of asparagine and glutamine set as dynamic and carbamidomethylations of cysteine set as static. N-glycosylation modifications were searched against the N-glycans mammalian database by Byonic™. The false discovery rate (FDR) was set to 1%. A focused database with decoys was created from the initial analysis and the focused database was used subsequently for all samples analysed. For samples that have mutations at a particular amino acid, the amino acid mutation was included as part of the modification parameters and set to dynamic.

Quantification of glycopeptides was based on peak area performed using the Minora feature detector and precursor ions quantifier nodes in Thermo Proteome Discoverer.

### Molecular dynamics simulations

The E protein dimer ectodomain (residues 1–395) was extracted from the experimental structure (PDB: 3J27) of the DENV2 NGC strain [68]. Amino acid mutations were introduced *in silico* using CHARMM-GUI to match the studied DENV2 D2Y98P strain [69]. Protein charges were assigned for neutral pH with charged termini. Based on the cell line, the predominant glycoforms at the N67-linked glycosylation site were as follows: i) HexNAc(2)Hex(6) for both C6/36 and Vero cells; and ii) HexNAc(4)Hex(5)NeuAc(2) in Huh-7 cells. For the N153-linked glycosylation site, the predominant glycoforms were: iii) HexNAc(2)Hex(3)Fuc(1) for C6/36; iv) HexNAc(5)Hex(5)Fuc(1) for Vero; and v) HexNAc(4)Hex(5)Fuc(1)NeuAc(1) in Huh-7. A total of five simulation constructs were created, including three simulations of DENV2 WT grown in C3/36, Vero, or Huh-7 with unique glycoforms at both N-linked glycosylation sites. Additionally, N153Q mutant simulations were conducted in C6/36 and Vero with an identical glycoform at N67, and in Huh-7. Each construct was placed in a dodecahedron box with an ∼18 nm edge length, containing around 120,000 TIP3P water molecules and 150 mM NaCl which also served to neutralize the system’s overall charge [70]. Energy minimization was performed using the steepest descent minimization algorithm with a 0.01 nm step size. Equilibration followed in the NVT and NPT ensembles for 5 ns each, with position restraints applied to protein alpha-carbons and glycan heavy atoms using a force constant 1000 kJ mol⁻¹ nm⁻². All systems were subjected to a production run of 500 ns in the NPT ensemble which involved weak position restraints on protein alpha-carbons with a force constant of 100 kJ mol^-1^ nm^-2^. All simulations utilized GROMACS 2018.3 with the CHARMM36m force field [71, 72]. To enhance glycan dynamics while retaining the protein structure, a temperature of 400 K was used. It was maintained using the velocity rescaling thermostat with an additional stochastic term using a time constant of 1 ps. Pressure was maintained isotropically at 1 atm using the Parrinello-Rahman barostat and a time constant of 5 ps [73]. Bonds involving hydrogens were constrained using the LINCS algorithm. Equations of motion were integrated using the leap-frog algorithm with a time step of 2 fs. Long-range electrostatic interactions were described using the particle mesh Ewald method [74]. The short-range electrostatics reals pace cut-off was 1.2 nm and the short-range van der Waals cut-off was also 1.2 nm. Periodic boundaries conditions were applied in all directions. All simulations snapshots were generated using VMD [75]. The glycan-protein contacts were calculated based on the 0.3 nm cutoff distance and were averaged over the entire 500 ns of trajectory. Each protein residue which interacted with glycans for over 10% of the total simulation time was highlighted. The 2D22 epitope on DENV2 dimer was taken from PDB:4UIH based on the 0.3 nm cutoff distance between any ectodomain residue and any Fab residue [32].

### Hydrogen Deuterium Exchange (HDX) mass spectrometry

DENV2 WT and N153Q mutant were grown in C6/36 cells and purified as described previously [76,77]. Deuterium exchange of purified E proteins solubilized in aqueous 100mM NaCl, 10 mM Tris-HCl pH 8.0 (NTE buffer) was initiated by incubation at 37°C in PBS buffer reconstituted in D_2_O (99.9%) resulting in a final D_2_O concentration of 90% as described previously [78]. Deuterium labeling was performed for 1 min for both WT and N153Q E proteins by quenching the exchange reaction by adding prechilled quench buffer. Quench buffers contain 1.5 M GdnHCl and 0.25 M Tris(2-carboxyethyl) phosphine-hydrochloride (TCEP-HCl) and after adding quench buffer the solutions were incubated at 4 °C on ice for 30 s followed by addition of titanium dioxide to precipitate envelope lipids. Precipitated envelope lipids were removed before injecting the sample for pepsin digestion using 0.22 μm centrifugal filters at 10,000 rpm for 1min.

Mass Spectrometry and peptide identification: Quenched samples were injected onto nanoUPLC HDX sample manager (Waters, Milford, MA). Injected samples were proteolyzed in online mode using pepsin immobilized Waters Enzymate column (2.1 × 30 mm) in 0.1% formic acid in water at a flow rate of 100 μl/min. The proteolyzed peptides were trapped in a 2.1 × 5 mm C18 trap (ACQUITY BEH C18 VanGuard Pre-column, 1.7 μm, Waters, Milford, MA). Elution of pepsin digested peptides was performed using acetonitrile (ACN) gradient of 8 to 40 % in 0.1 % formic acid at a flow rate of 40 μl min-1 into reverse phase column (ACQUITY UPLC BEH C18 Column, 1.0 × 100 mm, 1.7 μm, Waters) pumped by nanoACQUITY Binary Solvent Manager (Waters, Milford, MA). Peptides were ionized using electrospray ionization mode and sprayed onto SYNAPT G2-Si mass spectrometer (Waters, Milford, MA). HDMSE mode acquisition and measurement were employed. 200 fmol μl−1 of [Glu1]-fibrinopeptide B ([Glu1]-Fib) was injected at a flow rate of 5μl/min into mass spectrometer for calibration and lockspray correction.

Protein Lynx Global Server v3.0 (PLGS v3.0) was used to identify the peptides from undeuterated mass spectra (HDMS^E^). Search for peptide identification was performed on sequence database of D2Y98P strain with E, M and C proteins. No specific protease and variable N-linked glycosylation modifications were chosen in search parameters to carry out the sequence identification. Further, peptide identification parameters intensity of 2500 for product and precursor ions, minimum products per amino acids of 0.2 and a precursor ion mass tolerance of <10 ppm using DynamX v.3.0 (Waters, Milford, MA). Peptides present in at least two out of three undeuterated samples were retained. Reported deuterium exchange values are uncorrected for back exchange and all the reactions are performed in triplicates.

### Statistical analysis

Data were analyzed with GraphPad Prism 10.0. For statistical comparisons, the non-parametric Mann-Whitney U test was applied, while survival rates were compared using the Log-rank (Mantel-Cox) test. A p-value <0.05 (*) was considered statistically significant. Results were presented as mean ± standard deviation unless another method was specified in the figure legend.

## Acknowledgements

We thank Dr. Laurent Renia (A*STAR) for the kind gift of muMT mice. We also thank Dr. Ooi Eng Eong for sharing Huh-7 and pACYC177-CMV-HDVr-SV40PA plasmid.

## Author contributions

-Performed experiments: DHRT; JKM; CW; RVP; FI; ETXT; XNL; CVL; CHN

-Analysed data: DHRT; JKM; CW; WTCK; XNL; KRC; GSA; SA

-Provided reagents/inputs: IW; TNK; PJB; TLT; TSCB; PAM; YSL;

-Designed the experiments: DHRT; SA

-Wrote the manuscript: DHRT; SA

Donald Heng Rong Ting^1,2^, Jan Kazimierz Marzinek^3^, Corrine Wan^4^, Raghuvamsi V. Palur^5^, Fakhriedzwan Idris^1,2^, Eunice Tze Xin Tan^1,2^, Wei Teng Clara Koh^6^, Xin Ni Lim^6^, Tun-Linn Thein^7^, Cheer Vee Lim^1,2^, Chin Huan Ng^1,2^, Theresa S. C. Buckley^8^, Ian Walsh^4^, Paul A MacAry^2,9^, Yee-Sin Leo^7,10-13^, Ganesh S. Anand^8^, Kuan Rong Chan^6^, Terry Nguyen-Khuong^4^, Peter John Bond^3,14^, Sylvie Alonso^1,2*^

## Funding

This work was supported by the National Medical Research Council (NMRC) under grant # MOH-000087 allocated to SA. Work performed by PJB and JKM was supported by the National Research Foundation Competitive Research Programme (NRF-CRP27-2021-0003) and Bioinformatics Institute (A*STAR) core funds.

---

**Supplementary Table S1.** Deglycosylated E protein DENV2 mutants generated by site-directed mutagenesis. Mutation stability was assessed after three consecutive passages in C6/36 cell line.

| Position | Mutant | Amino acid substitution | Genetic stability |
| --- | --- | --- | --- |
| **N67** | N67A | Asn to Ala | No |
|  | N67D | Asn to Asp |  |
|  | N67K | Asn to Lys |  |
|  | N67L | Asn to Leu |  |
|  | N67S | Asn to Ser |  |
|  | N67T | Asn to Thr |  |
|  | N67Q | Asn to Gln |  |
|  | N67V | Asn to Val |  |
|  | T69A | Thr to Ala |  |
|  | T69D | Thr to Asp |  |
|  | T69K | Thr to Lys |  |
|  | T69Q | Thr to Gln |  |
|  | T69L | Thr to Leu |  |
|  | T69V | Thr to Val |  |
| **N153** | N153Q | Asn to Gln | Yes |

**Supplementary Table S2.**
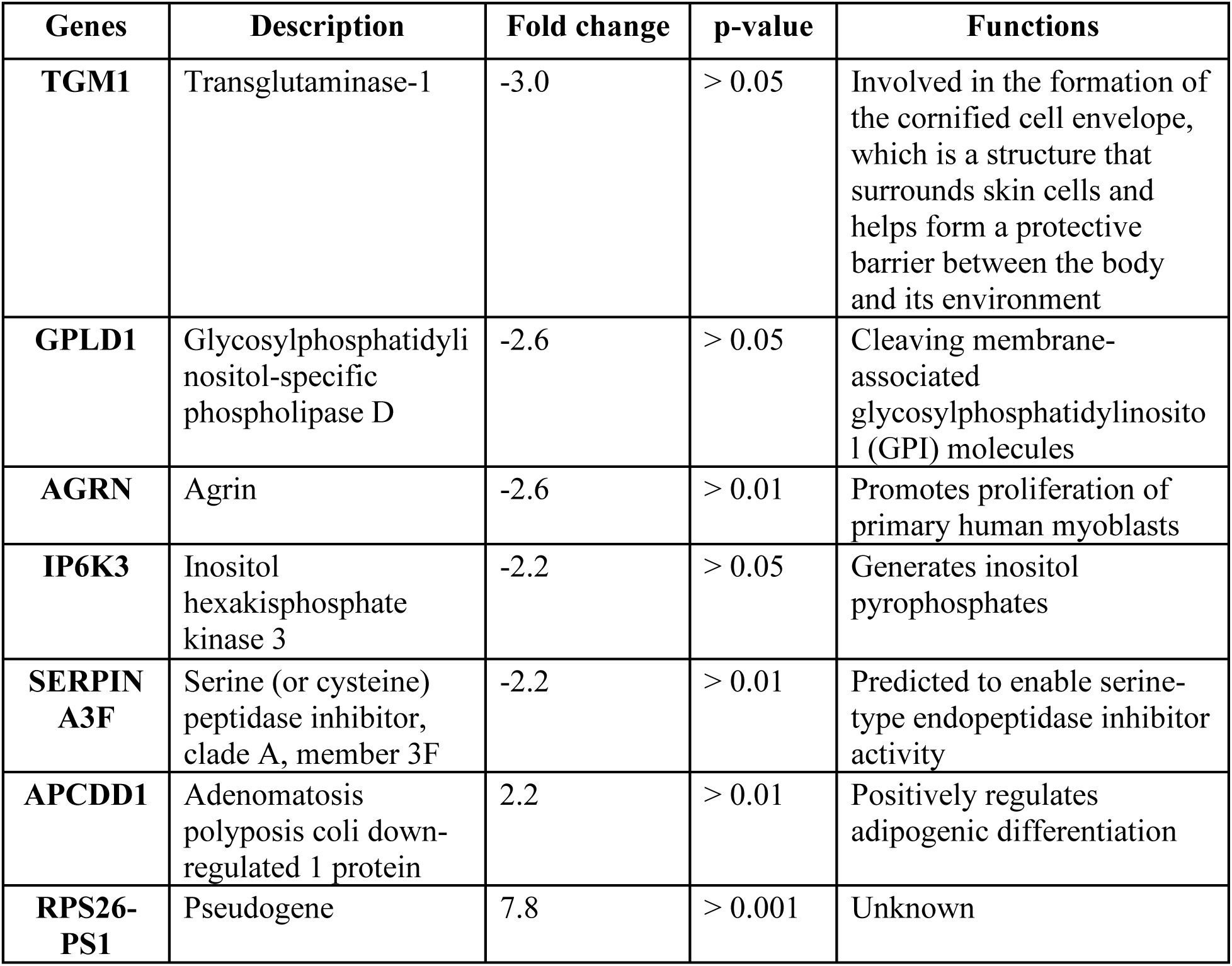
DEGs in N153Q mutant-infected versus WT-infected mice. Bulk RNAseq analysis was performed on white blood cells harvested at day 2 pi. from IFNAR^-/-^ mice infected with WT or N153Q mutant at 10^5^ PFU/mouse.

**Supplementary Table S3.** Amino acid shielded by N67- or N153-linked glycans in WT DENV2 grown in different cell lines. Amino acid residues that interacted with glycans for over 10% of the total simulation time in molecular dynamic simulations are indicated.

| **Cell line used to propagate virus** | **Amino acids shielded by N67-linked glycans** | **Amino acids shielded by N153-linked glycans** |
| --- | --- | --- |
| **C6/36** | EDII: 66-69, 83, 84, 89, 90, 118 | EDI: 2, 6, 148-158<br>EDII: 68-72, 99, 101-104, 246-248 |
| **Vero** | EDII: 66-69, 83, 84, 89, 90, 118 | EDI: 147-158, 176,<br>EDII: 68-73, 99, 101-105, 246, 247<br>EDII: 310 |
| **Huh-7** | EDII: 64, 66-73, 82-84, 89, 90, 118, 120, 122 | EDI: 46, 47, 149, 150, 152-158<br>EDII: 64-73, 77, 99, 101-105, 113, 202-204, 246-252, 271, 273-275, 277<br>EDIII: 310 |

**Supplementary Figure S1.**
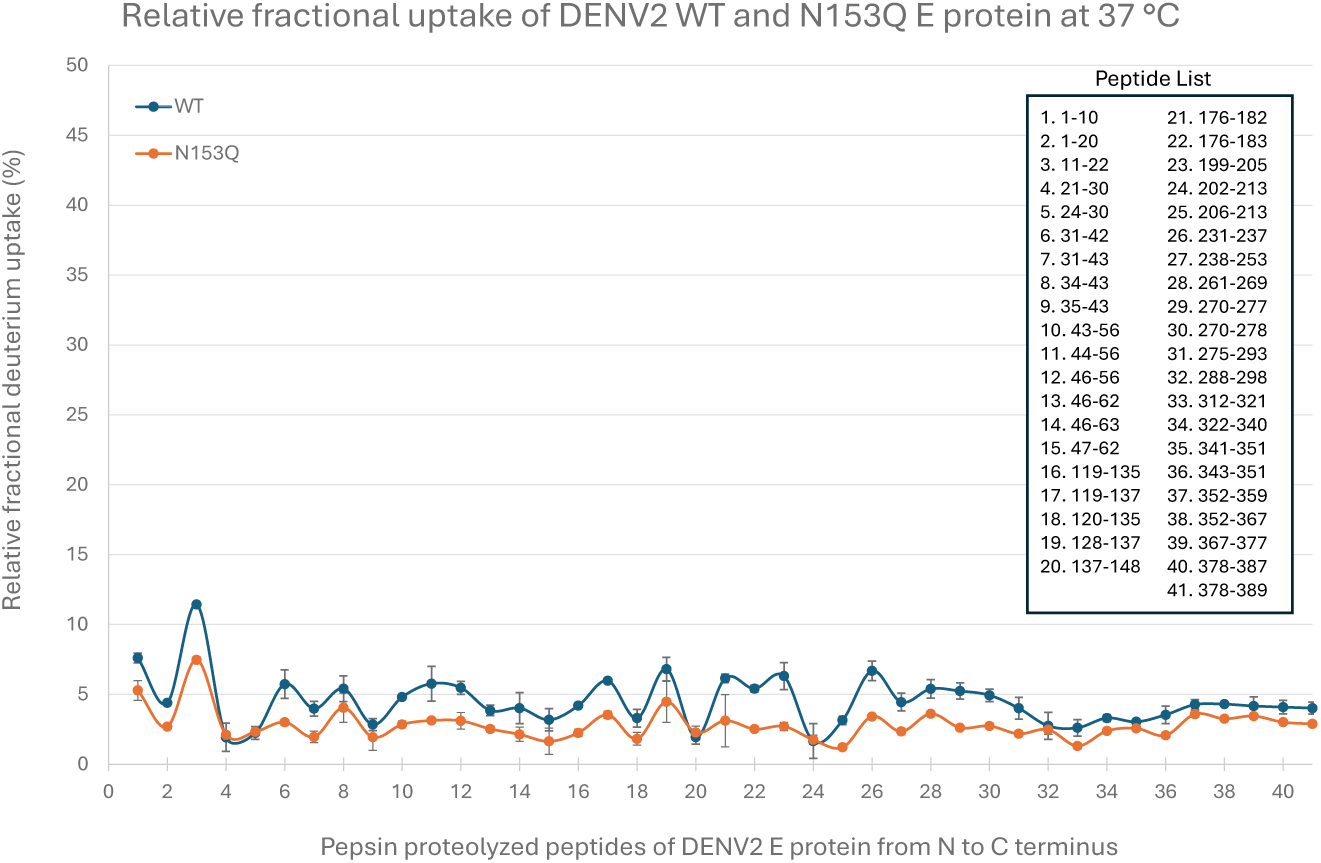
Deuterium uptake profiles of E protein from WT and N153Q DENV2. Relative deuterium fractional uptake plot showing the uptake for DENV2 E WT (Blue) and DENV2 E N153Q (Orange) at 37 °C at Dex=1 min. Relative fractional uptake is [(deuterium uptake in Da)/(total number of exchangeable backbone amide hydrogens)] * 100. Common pepsin fragment peptides between WT E and N153Q are represented by dots, listed from N to C terminus as listed. Error bars represent the relative standard deviation of deuterium exchange for each peptide.

**Supplementary Figure S2.**
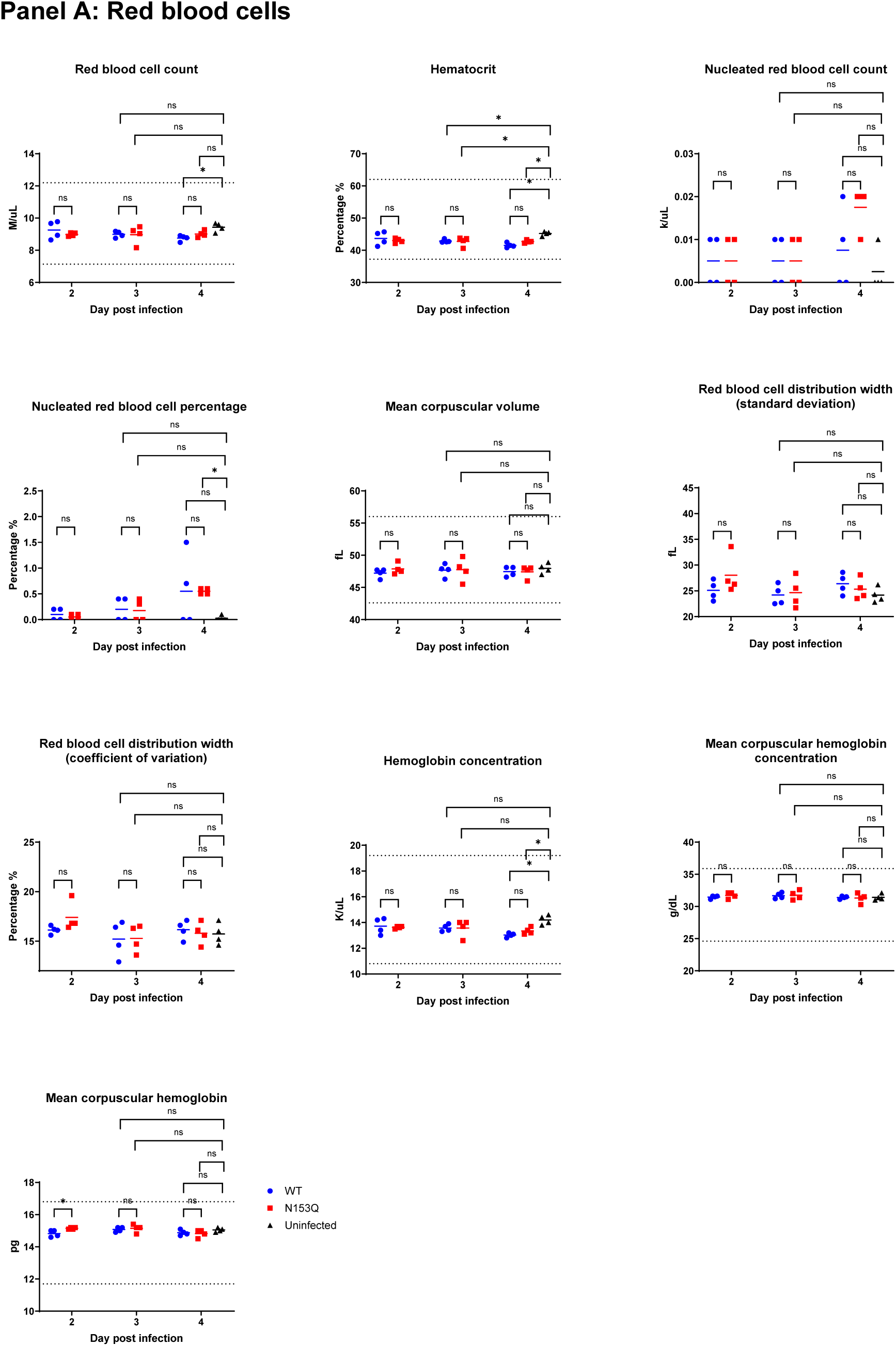
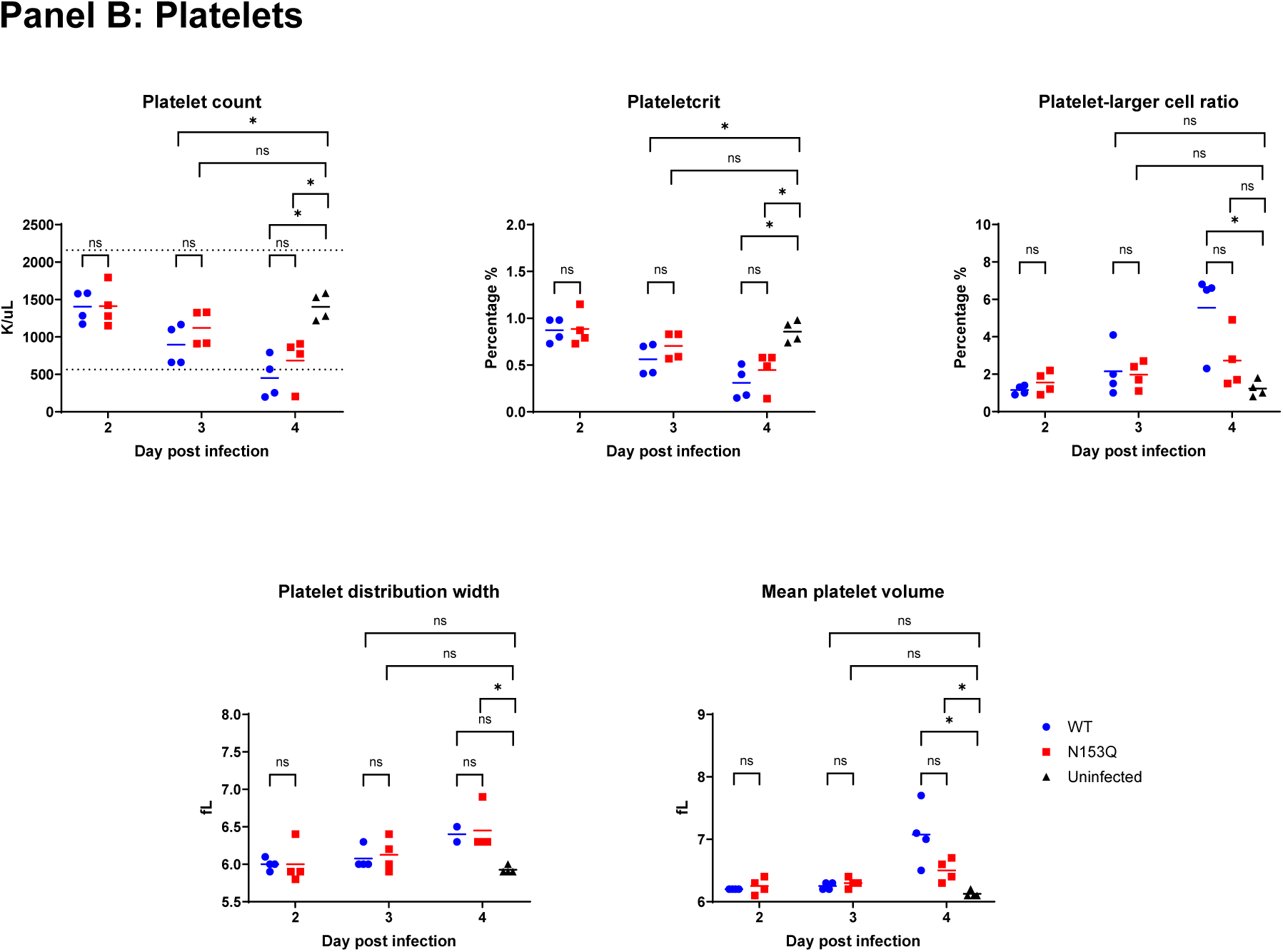
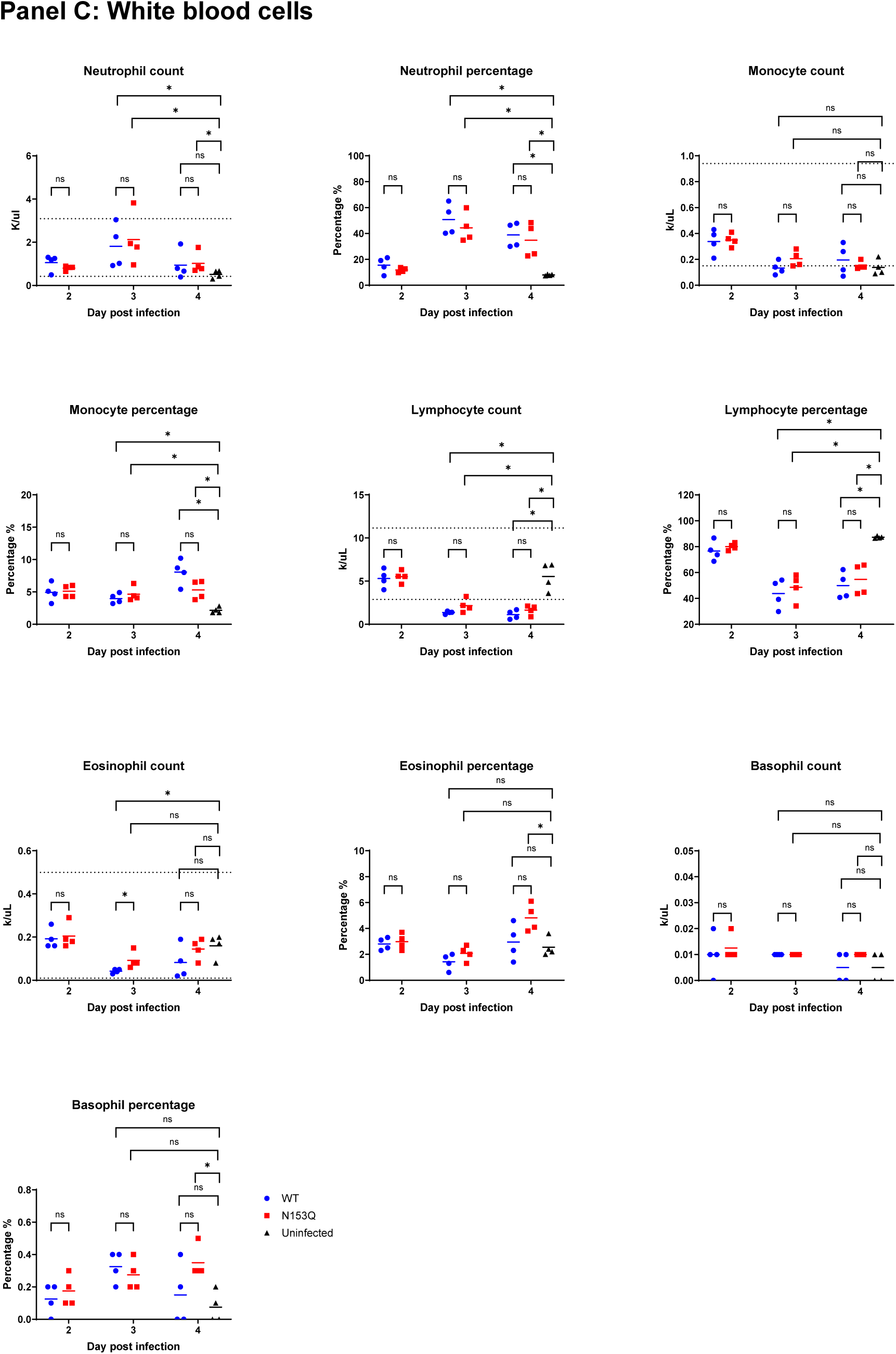
Blood parameters in IFNAR^-/-^ mice infected with WT and N153Q DENV2. 5-7 weeks old mice (n=4) were sc. infected with WT or N153Q at 10^5^ PFU/mouse. Blood was collected at the indicated time points to determine various cell counts as shown. Panel A: Red blood cells; Panel B: Platelets; Panel C: White blood cells. K/uL, thousands per cubic millilitre; M/uL, millions per cubic millilitre; fL, femtoliters; g/dL, grams per decilitre; pg, picograms. Dotted lines indicated the lower and upper limits of reference range. Statistical analysis was performed using nonparametric Mann–Whitney U test. *, p < 0.05; ns, not significant.

**Supplementary Figure S3.**
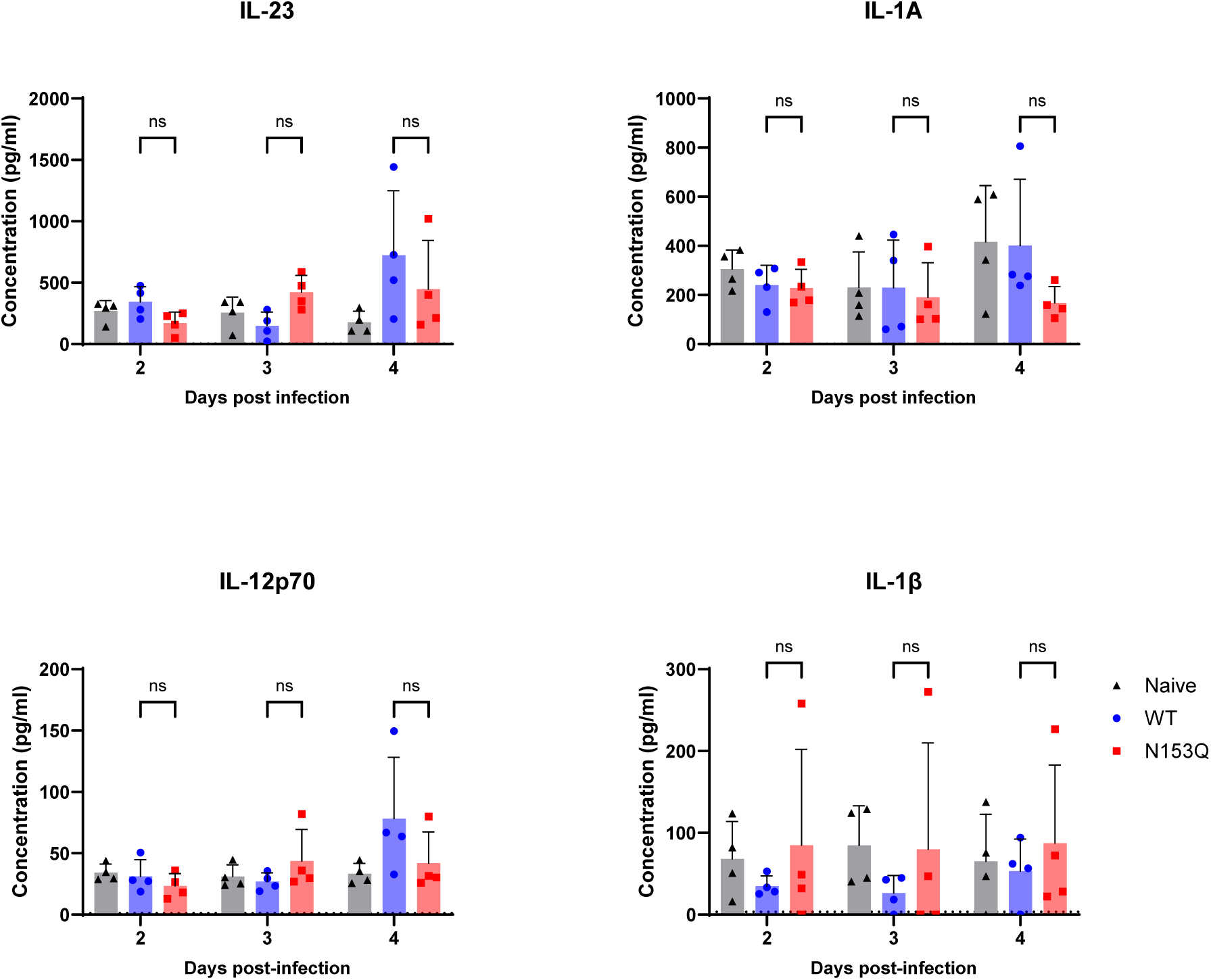
Systemic inflammatory cytokines levels in IFNAR^-/-^ mice infected with WT or N153Q DENV2. 5-7 weeks old mice (n=4) were sc. infected with WT or N153Q DENV2 at 10^5^ PFU/mouse. Systemic levels of IL-23, IL-1A, IL-12p70 and IL-1β were measured by multiplex assay. Data were expressed as the mean +/- SD. Data shown were representative of at least two independent experiments. Statistical analysis was performed using nonparametric Mann–Whitney U test. *, p < 0.05; ns, not significant.

**Supplementary Figure S4.**
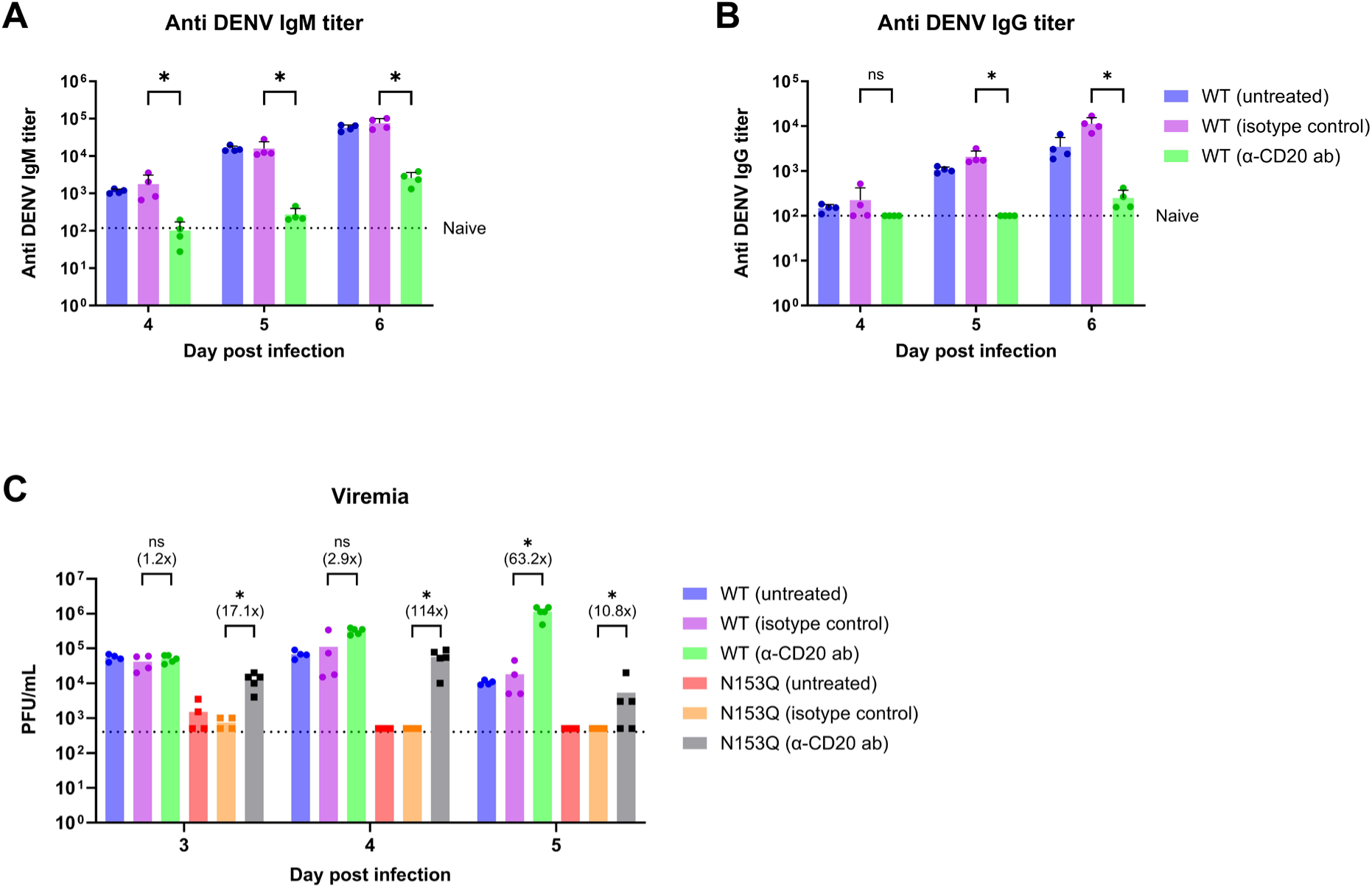
N153Q mutant infection in B cell-depleted IFNAR^-/-^ mice. 5-7 weeks old mice (n=4) were injected ip. with 100µg of anti-CD20 mAb or isotype control mAb two days prior to sc. infection with WT or N153Q DENV2 at 10^5^ PFU/mouse. Systemic DENV-specific IgM (A) and IgG (B) antibody titers in WT-infected mice were measured by ELISA at the indicated time points pi. WT virus grown in Vero cells was used as coating antigen. Dotted line indicates the mean Ab titers detected in naïve mice. (C) Viremia titers were measured by plaque assay. Dotted line indicates the detection limit of the assay. Data are representative of at least two independent experiments. Statistical analysis was performed using nonparametric Mann–Whitney U test. *, p < 0.05; ns, not significant; differences in fold change were indicated in bracket.

**Supplementary Figure S5.**
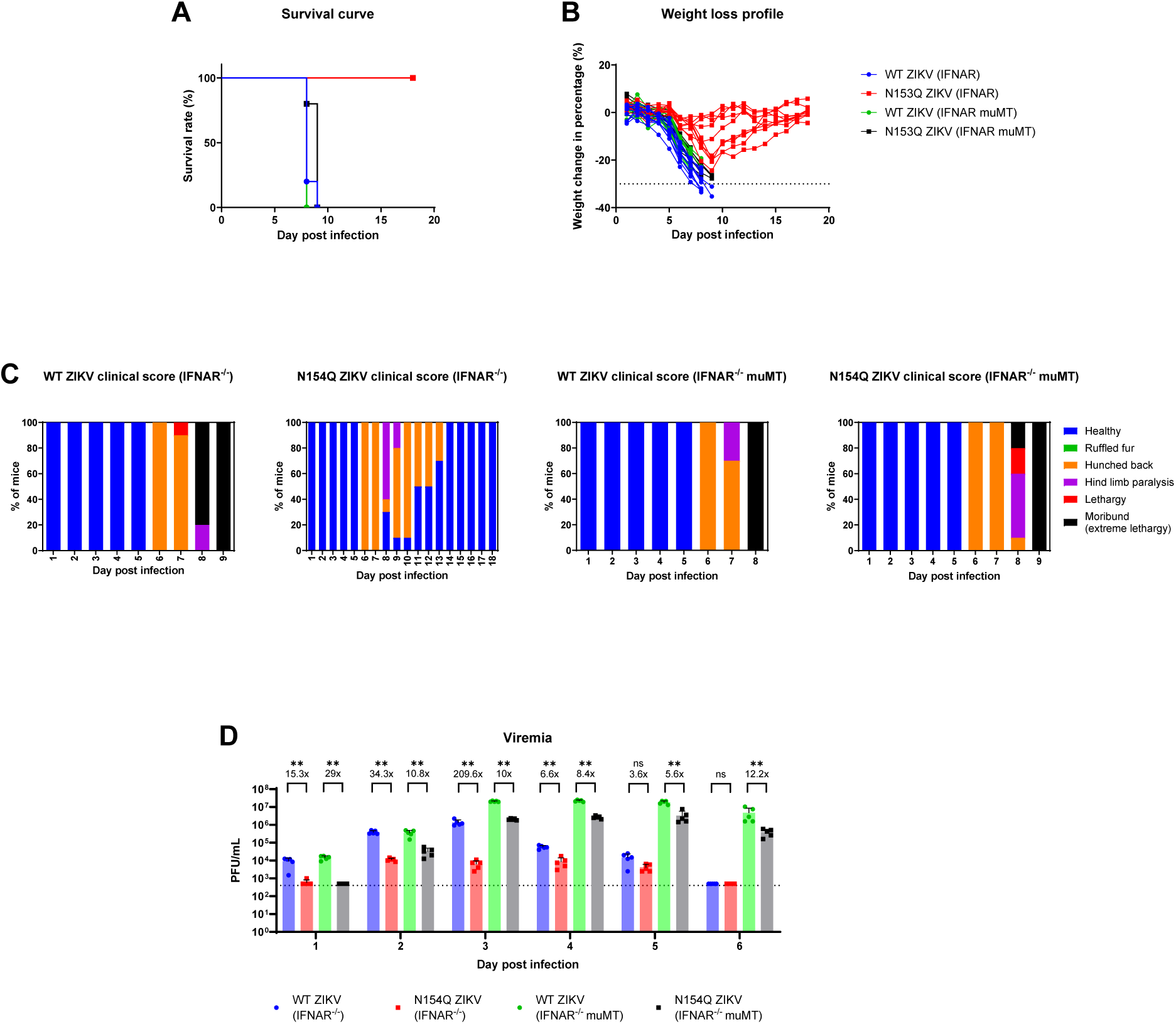
N154Q ZIKV mutant infection profile in IFNAR^-/-^ and muMT IFNAR^-/-^ mice. 5-7 weeks old mice (n=7/8) were sc. infected with WT or N154Q ZIKV at 10^6^ PFU/mouse. (A) Kaplan-Meier survival curve. (B) Body weight loss profile. (C) Clinical scores. Mice that lost 20% of their body weight measured at the start of infection and/or that were moribund were promptly euthanized. (D) Viremia titers (n=5) were measured by plaque assay. Dotted line indicates the detection limit of the assay. Data are representative of at least two independent experiments. Statistical analysis was performed using nonparametric Mann–Whitney U test. *, p < 0.05; **, p < 0.01; ns, not significant.

**Supplementary Figure S6.**
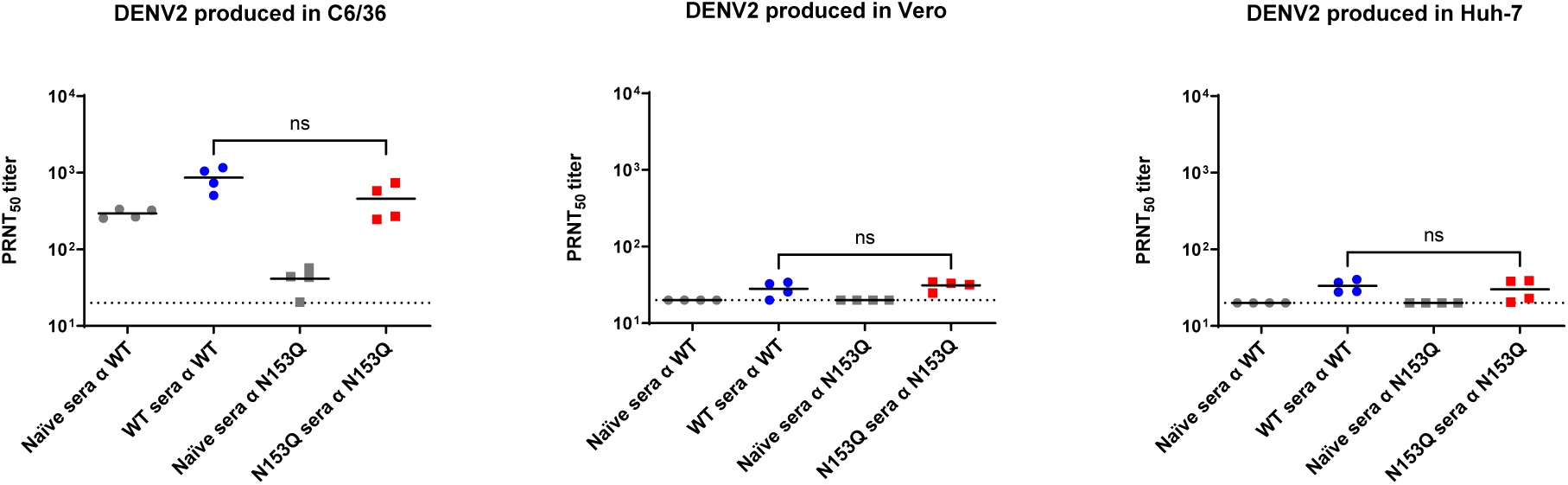
Sensitivity of DENV2 to mouse sera. PRNT_50_ titers of mouse sera were determined against WT and N153Q virus produced in C6/36, Vero and Huh-7 cell lines. 5-7 weeks old IFNAR^-/-^ mice (n=4) were sc. infected with WT or N153Q mutant at 10^5^ PFU/mouse. Serum was obtained at day 5 pi. and heat-inactivated and neutralization assay was conducted in homologous fashion. Dotted line indicates the detection limit of the assay. Statistical analysis was performed using student t-test.

**Supplementary Figure S7.**
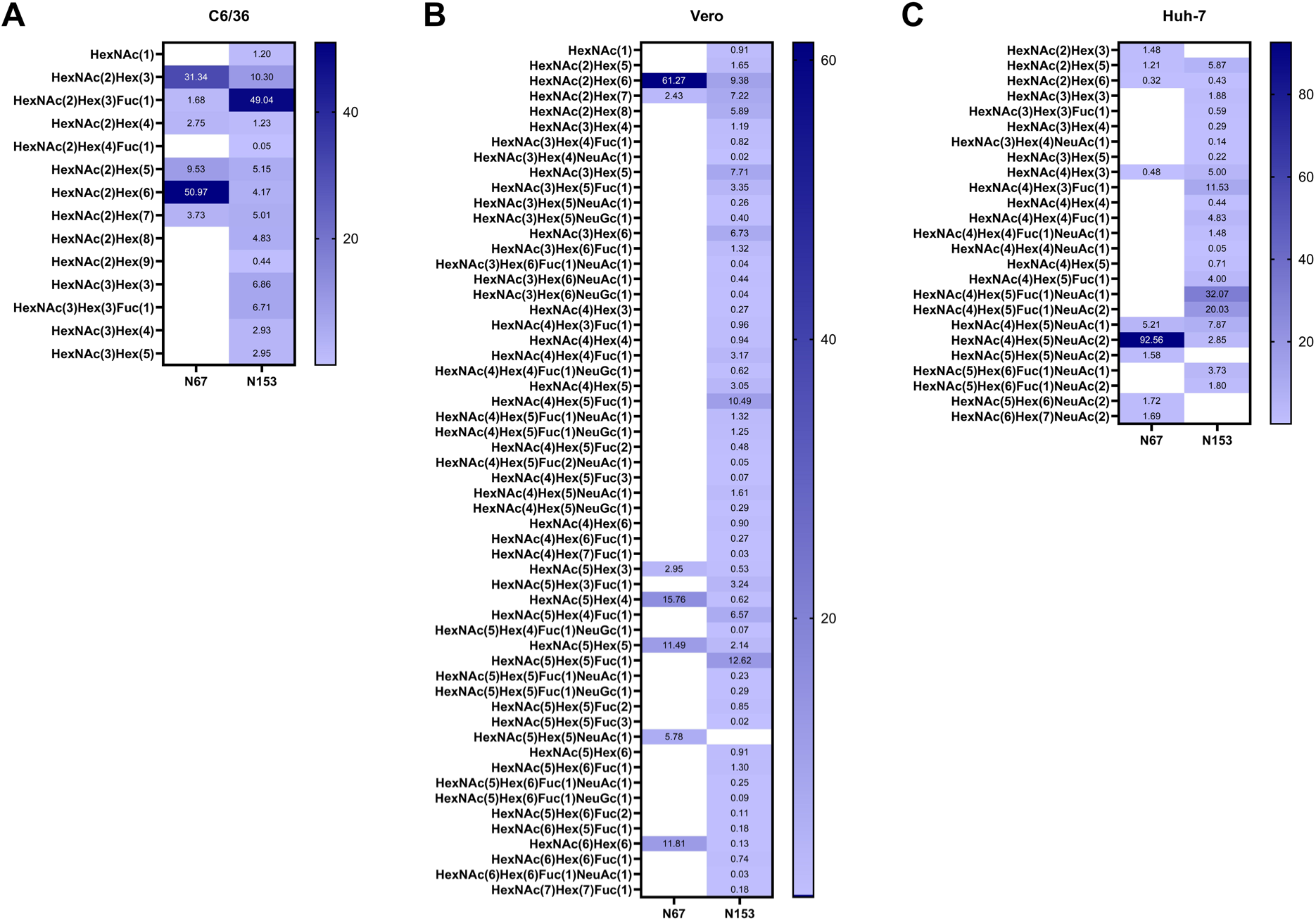
Glycoforms detected at N67 and N153 on E protein. WT DENV2 was produced in C6/36, Vero or Huh-7 cell lines. Virus particles in the culture supernatant were harvested and processed for glycoproteomics analysis. Data were expressed as the percentage of each glycoform found at N67 and N153. Data represent mean value of two independent biological replicates. Hex, Hexose; HexNAc, N-Acetylhexosamine; Fuc, fucose. NeuAc, N-acetylneuraminic acid; NeuGc, N-Glycolylneuraminic acid.

## Notes

### Competing Interest Statement

The authors have declared no competing interest.

## References

1. Brady, O.J., et al., Refining the global spatial limits of dengue virus transmission by evidence-based consensus. PLoS Negl Trop Dis, 2012. 6(8): p. e1760.

2. Bhatt, S., et al., The global distribution and burden of dengue. Nature, 2013. 496(7446): p. 504–7.

3. Messina, J.P., et al., The current and future global distribution and population at risk of dengue. Nat Microbiol, 2019. 4(9): p. 1508–1515.

4. World Health Organization, Dengue: guidelines for diagnosis, treatment, prevention and control. 2009: World Health Organization.

5. Burke, D.S., et al., A prospective study of dengue infections in Bangkok. Am J Trop Med Hyg, 1988. 38(1): p. 172–80.

6. Guzman, M.G., et al., Enhanced severity of secondary dengue-2 infections: death rates in 1981 and 1997 Cuban outbreaks. Rev Panam Salud Publica, 2002. 11(4): p. 223–7.

7. Kaptein, S.J.F., et al., A pan-serotype dengue virus inhibitor targeting the NS3-NS4B interaction. Nature, 2021. 598(7881): p. 504–509.

8. Goethals, O., et al., Blocking NS3-NS4B interaction inhibits dengue virus in non-human primates. Nature, 2023. 615(7953): p. 678–686.

9. Ackaert, O., et al., Safety, Tolerability, and Pharmacokinetics of JNJ-1802, a Pan-serotype Dengue Direct Antiviral Small Molecule, in a Phase 1, Double-Blind, Randomized, Dose-Escalation Study in Healthy Volunteers. Clin Infect Dis, 2023. 77(6): p. 857–865.

10. Waickman, A.T., et al., Biologics for dengue prevention: up-to-date. Expert Opin Biol Ther, 2023. 23(1): p. 73–87.

11. Sabchareon, A., et al., Protective efficacy of the recombinant, live-attenuated, CYD tetravalent dengue vaccine in Thai schoolchildren: a randomised, controlled phase 2b trial. Lancet, 2012. 380(9853): p. 1559–67.

12. Capeding, M.R., et al., Clinical efficacy and safety of a novel tetravalent dengue vaccine in healthy children in Asia: a phase 3, randomised, observer-masked, placebo-controlled trial. Lancet, 2014. 384(9951): p. 1358–65.

13. Villar, L., et al., Efficacy of a tetravalent dengue vaccine in children in Latin America. N Engl J Med, 2015. 372(2): p. 113–23.

14. Biswal, S., et al., Efficacy of a tetravalent dengue vaccine in healthy children aged 4-16 years: a randomised, placebo-controlled, phase 3 trial. Lancet, 2020. 395(10234): p. 1423–1433.

15. Halstead, S.B., Dengvaxia sensitizes seronegatives to vaccine enhanced disease regardless of age. Vaccine, 2017. 35(47): p. 6355–6358.

16. Postler, T.S., et al., Renaming of the genus Flavivirus to Orthoflavivirus and extension of binomial species names within the family Flaviviridae. Arch Virol, 2023. 168(9): p. 224.

17. Idris, F., D.H.R. Ting, and S. Alonso, An update on dengue vaccine development, challenges, and future perspectives. Expert Opin Drug Discov, 2021. 16(1): p. 47–58.

18. Hacker, K., L. White, and A.M. de Silva, N-linked glycans on dengue viruses grown in mammalian and insect cells. J Gen Virol, 2009. 90(Pt 9): p. 2097–106.

19. Johnson, A.J., F. Guirakhoo, and J.T. Roehrig, The envelope glycoproteins of dengue 1 and dengue 2 viruses grown in mosquito cells differ in their utilization of potential glycosylation sites. Virology, 1994. 203(2): p. 241–9.

20. Lei, Y., et al., Characterization of N-Glycan Structures on the Surface of Mature Dengue 2 Virus Derived from Insect Cells. PLoS One, 2015. 10(7): p. e0132122.

21. Dubayle, J., et al., Site-specific characterization of envelope protein N-glycosylation on Sanofi Pasteur’s tetravalent CYD dengue vaccine. Vaccine, 2015. 33(11): p. 1360–8.

22. Alen, M.M., et al., Crucial role of the N-glycans on the viral E-envelope glycoprotein in DC-SIGN-mediated dengue virus infection. Antiviral Res, 2012. 96(3): p. 280–7.

23. Miller, J.L., et al., The mannose receptor mediates dengue virus infection of macrophages. PLoS Pathog, 2008. 4(2): p. e17.

24. Chen, S.T., et al., CLEC5A is critical for dengue-virus-induced lethal disease. Nature, 2008. 453(7195): p. 672–6.

25. Pokidysheva, E., et al., Cryo-EM reconstruction of dengue virus in complex with the carbohydrate recognition domain of DC-SIGN. Cell, 2006. 124(3): p. 485–93.

26. Bryant, J.E., et al., Glycosylation of the dengue 2 virus E protein at N67 is critical for virus growth in vitro but not for growth in intrathoracically inoculated Aedes aegypti mosquitoes. Virology, 2007. 366(2): p. 415–23.

27. Lee, E., et al., Both E protein glycans adversely affect dengue virus infectivity but are beneficial for virion release. J Virol, 2010. 84(10): p. 5171–80.

28. Mondotte, J.A., et al., Essential role of dengue virus envelope protein N glycosylation at asparagine-67 during viral propagation. J Virol, 2007. 81(13): p. 7136–48.

29. Lee, E., R.C. Weir, and L. Dalgarno, Changes in the dengue virus major envelope protein on passaging and their localization on the three-dimensional structure of the protein. Virology, 1997. 232(2): p. 281–90.

30. Guirakhoo, F., et al., Selection and partial characterization of dengue 2 virus mutants that induce fusion at elevated pH. Virology, 1993. 194(1): p. 219–23.

31. Dejnirattisai, W., et al., A new class of highly potent, broadly neutralizing antibodies isolated from viremic patients infected with dengue virus. Nat Immunol, 2015. 16(2): p. 170–177.

32. Fibriansah, G., et al., DENGUE VIRUS. Cryo-EM structure of an antibody that neutralizes dengue virus type 2 by locking E protein dimers. Science, 2015. 349(6243): p. 88–91.

33. Sharma, A., et al., The epitope arrangement on flavivirus particles contributes to Mab C10’s extraordinary neutralization breadth across Zika and dengue viruses. Cell, 2021. 184(25): p. 6052–6066 e18.

34. Zhang, S., et al., Neutralization mechanism of a highly potent antibody against Zika virus. Nat Commun, 2016. 7: p. 13679.

35. Fontes-Garfias, C.R., et al., Functional Analysis of Glycosylation of Zika Virus Envelope Protein. Cell Rep, 2017. 21(5): p. 1180–1190.

36. Annamalai, A.S., et al., Zika Virus Encoding Nonglycosylated Envelope Protein Is Attenuated and Defective in Neuroinvasion. J Virol, 2017. 91(23).

37. Carbaugh, D.L., R.S. Baric, and H.M. Lazear, Envelope Protein Glycosylation Mediates Zika Virus Pathogenesis. J Virol, 2019. 93(12).

38. Liang, J.J., M.W. Chou, and Y.L. Lin, DC-SIGN Binding Contributed by an Extra N-Linked Glycosylation on Japanese Encephalitis Virus Envelope Protein Reduces the Ability of Viral Brain Invasion. Front Cell Infect Microbiol, 2018. 8: p. 239.

39. Beasley, D.W., et al., Envelope protein glycosylation status influences mouse neuroinvasion phenotype of genetic lineage 1 West Nile virus strains. J Virol, 2005. 79(13): p. 8339–47.

40. Yenamandra, S.P., et al., Evolution, heterogeneity and global dispersal of cosmopolitan genotype of Dengue virus type 2. Sci Rep, 2021. 11(1): p. 13496.

41. Graf, T., et al., Multiple introductions and country-wide spread of DENV-2 genotype II (Cosmopolitan) in Brazil. Virus Evol, 2023. 9(2): p. vead059.

42. Cheang, Y.Z.N., et al., In vitro and in vivo efficacy of Metformin against dengue. Antiviral Res, 2021. 195: p. 105186.

43. Zompi, S., et al., Protection from secondary dengue virus infection in a mouse model reveals the role of serotype cross-reactive B and T cells. J Immunol, 2012. 188(1): p. 404–16.

44. Haas, K.M., et al., Protective and pathogenic roles for B cells during systemic autoimmunity in NZB/W F1 mice. J Immunol, 2010. 184(9): p. 4789–800.

45. Rouvinski, A., et al., Recognition determinants of broadly neutralizing human antibodies against dengue viruses. Nature, 2015. 520(7545): p. 109–13.

46. Lambert, G.S. and C. Upadhyay, HIV-1 Envelope Glycosylation and the Signal Peptide. Vaccines (Basel), 2021. 9(2).

47. Seabright, G.E., et al., Protein and Glycan Mimicry in HIV Vaccine Design. J Mol Biol, 2019. 431(12): p. 2223–2247.

48. Pralow, A., et al., Comprehensive N-glycosylation analysis of the influenza A virus proteins HA and NA from adherent and suspension MDCK cells. FEBS J, 2021. 288(16): p. 4869–4891.

49. Gong, S. and R.M. Ruprecht, Immunoglobulin M: An Ancient Antiviral Weapon - Rediscovered. Front Immunol, 2020. 11: p. 1943.

50. Kubagawa, H., et al., Functional Roles of the IgM Fc Receptor in the Immune System. Front Immunol, 2019. 10: p. 945.

51. Kubagawa, H., et al., Differences between Human and Mouse IgM Fc Receptor (FcmicroR). Int J Mol Sci, 2021. 22(13).

52. Kubli, S.P., et al., Reply to: Questioning whether the IgM Fc receptor (FcmuR) is expressed by innate immune cells. Nat Commun, 2022. 13(1): p. 3950.

53. Skopnik, C.M., et al., Questioning whether IgM Fc receptor (FcmicroR) is expressed by innate immune cells. Nat Commun, 2022. 13(1): p. 3951.

54. Shibuya, A., et al., Fc alpha/mu receptor mediates endocytosis of IgM-coated microbes. Nat Immunol, 2000. 1(5): p. 441–6.

55. Fuchs, A., et al., Direct complement restriction of flavivirus infection requires glycan recognition by mannose-binding lectin. Cell Host Microbe, 2010. 8(2): p. 186–95.

56. Johnson, J.B., G.A. Capraro, and G.D. Parks, Differential mechanisms of complement-mediated neutralization of the closely related paramyxoviruses simian virus 5 and mumps virus. Virology, 2008. 376(1): p. 112–23.

57. Nakamura, M., et al., Complement-dependent virolysis of HIV-1 with monoclonal antibody NM-01. AIDS Res Hum Retroviruses, 1993. 9(7): p. 619–26.

58. Huber, M., et al., Complement lysis activity in autologous plasma is associated with lower viral loads during the acute phase of HIV-1 infection. PLoS Med, 2006. 3(11): p. e441.

59. Khoo, U.S., et al., DC-SIGN and L-SIGN: the SIGNs for infection. J Mol Med (Berl), 2008. 86(8): p. 861–74.

60. Amraei, R., et al., CD209L/L-SIGN and CD209/DC-SIGN Act as Receptors for SARS-CoV-2. ACS Cent Sci, 2021. 7(7): p. 1156–1165.

61. van der Zande, H.J.P., et al., The Mannose Receptor: From Endocytic Receptor and Biomarker to Regulator of (Meta)Inflammation. Front Immunol, 2021. 12: p. 765034.

62. Leo, Y.S., et al., Utility of warning signs in guiding admission and predicting severe disease in adult dengue. BMC Infect Dis, 2013. 13: p. 498.

63. Schreiber, M.J., et al., Genomic epidemiology of a dengue virus epidemic in urban Singapore. J Virol, 2009. 83(9): p. 4163–73.

64. Grant, D., et al., A single amino acid in nonstructural protein NS4B confers virulence to dengue virus in AG129 mice through enhancement of viral RNA synthesis. J Virol, 2011. 85(15): p. 7775–87.

65. Gavor, E., et al., Structure of Aedes aegypti procarboxypeptidase B1 and its binding with Dengue virus for controlling infection. Life Sci Alliance, 2022. 5(1).

66. Koh, C.W.T., et al., STAGEs: A web-based tool that integrates data visualization and pathway enrichment analysis for gene expression studies. Sci Rep, 2023. 13(1): p. 7135.

67. Hughes, C.S., et al., Single-pot, solid-phase-enhanced sample preparation for proteomics experiments. Nat Protoc, 2019. 14(1): p. 68–85.

68. Zhang, X., et al., Cryo-EM structure of the mature dengue virus at 3.5-A resolution. Nat Struct Mol Biol, 2013. 20(1): p. 105–10.

69. Jo, S., et al., CHARMM-GUI: a web-based graphical user interface for CHARMM. J Comput Chem, 2008. 29(11): p. 1859–65.

70. Jorgensen, W.L., et al., Comparison of simple potential functions for simulating liquid water. The Journal of Chemical Physics, 1983. 79(2): p. 926–935.

71. Van Der Spoel, D., et al., GROMACS: fast, flexible, and free. J Comput Chem, 2005. 26(16): p. 1701–18.

72. Huang, J., et al., CHARMM36m: an improved force field for folded and intrinsically disordered proteins. Nat Methods, 2017. 14(1): p. 71–73.

73. Parrinello, M. and A. Rahman, Polymorphic transitions in single crystals: A new molecular dynamics method. Journal of Applied Physics, 1981. 52(12): p. 7182–7190.

74. Essmann, U., et al., A smooth particle mesh Ewald method. The Journal of Chemical Physics, 1995. 103(19): p. 8577–8593.

75. Humphrey, W., A. Dalke, and K. Schulten, VMD: visual molecular dynamics. J Mol Graph, 1996. 14(1): p. 33–8, 27-8.

76. Fibriansah, G., et al. Structural changes in dengue virus when exposed to a temperature of 37 degrees C. Journal of Virology, 2013. 87: p. 7585–7592.

77. Kuhn, R. J., et al. Structure of dengue virus: Implications for flavivirus organization, maturation, and fusion. Cell 2002. 108: p.717–725.

78. Lim, X.X., et al. Conformational changes in intact dengue virus reveal serotype-specific expansion. Nature Communications, 2017. 8:14339.

